# A new hypothesis-testing model for phyllochron based on a stochastic process - application to analysis of genetic and environment effects in maize

**DOI:** 10.1101/2021.01.11.426247

**Authors:** S. Plancade, E. Marchadier, S. Huet, A. Ressayre, C. Noûs, C. Dillmann

## Abstract

The times between appearance of successive leaves or phyllochron characterize the vegetative development of annual plants. Hypothesis testing models, which enables to compare phyllochron between genetic groups or conditions, are usually based on regression of thermal time on the number of leaves, most of the time assuming a constant leaf appearance rate. However these models are both statistically biased and inappropriate in terms of modelling. We propose a stochastic process model in which the emergence of new leaves is considered as successive time-to-events, which provides a flexible and more accurate modelling as well as unbiased testing procedures. The model was applied on an original maize dataset collected in fields for three years on plants originating from two divergent selection experiments for flowering time conducted in two maize inbred lines. We showed that the main differences in phyllochron were not observed between selection populations (Early or Late), but rather between ancestral lines, years of experimentation, and leaf ranks. Our results highlight a strong departure from the assumption of a constant leaf appearance rate in one year that could be related to climate variations, even if the impact of each climatic variables individually was not clearly elucidated.

## 1 Introduction

In annual plant species, growth and development are critical to determine the time to completion of the life-cycle (from germination to seed maturation), that impacts biomass production. Plant development is characterized by the repeated production and growth of different organs that usually remain hidden and protected for a while in older tissues. Because it does not require destructive measurement, the time between leaf appearance, or phyllochron, is widely used to globally gauge the pace of plant development. Because plant growth rate varies with temperature, a transformation of the time-scale is usually performed through the use of accumulated thermal time (ATT) measured in degree-days (Parent and Tardieu, 2012). The objective of the transformation is to make plant growth rates constant in thermal time (van Straalen, 1983), thereby allowing temperature independent model to be used once the time has been transformed appropriately to integrate temperature.

Models for plant growth and development can be developed either to predict crop production in response to environmental factors and management practices or to investigate and test for differences in patterns of growth due to either environmental and/or genetic effects. The first category of models simulate crop development either at the field level using compartment models or at the plant level allowing for the description of plant architecture: (i) Compartmental models are based on differential rate equations that simulate how the plant community responds to the environment at the field scale (e.g. APSIM (Wang et al., 2002; Brown et al., 2018; Jamieson et al., 1998; He et al., 2012)); or (ii) Functional Structural Plant models focus on the development of organs of individual plants allowing studies of the interaction of plants within a field (Vidal and Andrieu, 2020; Vidal et al., 2021; Soualiou et al., 2021). Models (i) and (ii) are developed for prediction purpose, e.g. to predict biomass production of different genotypes in various climate scenarios, sowing densities, management practices. Another kind of models is required to compare growth rate or development transitions between classes (e.g. conditions or genotypic groups): Hypothesis testing models.

Regarding phyllochron analysis based on repeated measurements on plants from several classes (or conditions), the mostly used hypothesis testing model is the *linear phyllochron* which assumes a constant leaf appearance rate, inferred by linear regression of times of measurements (in ATT) on the numbers of leaves, separately on each plant (Padilla and Otegui, 2005; van Esbroeck et al., 2008; Correia et al., 2016). Thus, the phyllochron of each plant is summarised by a single value, and the difference of phyllochron between conditions can then be tested by a parametric or non-parametric univariate test (F-test, Mann-Whitney…). However this approach is simultaneously statistically biased and inapropriate in terms of modelling. First, the error of estimation of the single-plant phyllochron coefficient is ignored; Notably, the testing power is the same whatever the number of monitoring time points which is intuitively and statistically flawed. Secondly, the response of growth parameters to within-season variations of environmental variables has already been reported (Yu and Goh, 2019), as well as the independence of growth parameters between leaf rank (Chenu et al., 2008) which questions the assumption of phyllochron constancy. Some phyllochron models with more flexible functions of ATT have been proposed, in particular bi- or tri-linear regression for rice (Clerget and Bueno, 2013) or splines for wheat (Baumont et al., 2019). Even if they can accurately fit the phyllochron variations, these models are inadequate for hypothesis testing since the variations around the average curve are inappropriately modelled. Indeed, regression models implicitely assume that the phyllochron process (i.e. the curve of the number of leaves over time) is the sum of a general trend and of independent centered random variations. But simulating a phyllochron process from a regression model would generate non-increasing curves (while the number of leaves should increase over time), since centered random variations can be either negative or positive. More generally, the assumption of independent random variations is unrealistic, since the phyllochron process is by essence auto-correlated i.e. the number of leaves at time *t′* > *t* depends on the number of leaves at time *t*. This phenomenon results in a higher variability of leaf appearance time as the leaf rank increase (see e.g. Figure 1 in Millet et al. (2019)). We can mention an alternative which circumvents the interval censoring i.e. the fact that leaf appearance times are only known to lie in a certain time interval between two observation points: Zhu et al. (2014) consider that a leaf appears midway between the last observation when leaf was not observed and the first observation when it was, in a design with two observations per week. This approximation amounts to assume that the whole phyllochron process is observed, but it is biased unless interval between monitoring times are negligible with respect to interval between appearance of successive leaves.

**Figure 1:**
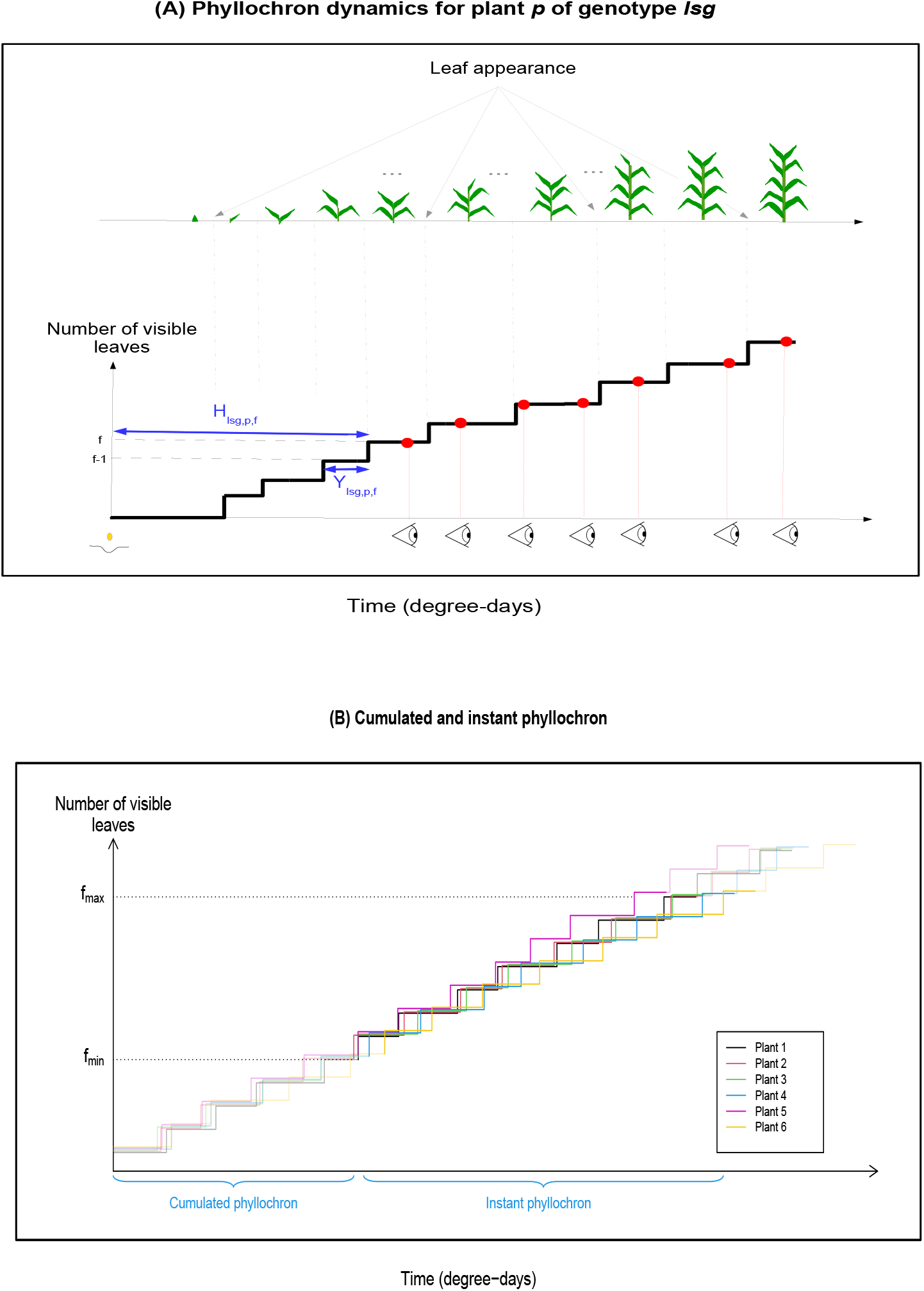
Illustration of the phyllochron process. (A) Phyllochron dynamics or process for a plant *p* belonging to genotype *lsg*. The step curve corresponds to the number of visible leaves over time; the interval between steps corresponds to the time interval between successive leaves denoted *Y_lsg,p,f_*; the time between sowing and appearance of leaf *f* is denoted *H_lsg,p,f_*, and is equal to the sum of the time intervals between leaves until leaves of rank smaller than f. Red bullets corresponds to the monitoring times, at which the number of leaves is recorded. (B) Simulated phyllochron dynamics for six plants from the same genotype (one color by plant). For each plant, the light-colored curve corresponds to the whole phyllochron i.e. to the process of appearance of all leaves, while the colorful curve corresponds to the phyllochron on the restricted range of leaf ranks [*f*_min_, *f*_max_] imposed by experimental constraints. The first modelled leaf *f*_min_ determines the *cumulated phyllochron* which corresponds to the time between sowing and appearance of leaf *f*_min_ and the *instant phyllochron* which corresponds to all intervals between successive leaves on the range [*f*_min_, *f*_max_].

In the context of a unique time-to-event, the statistical flaws of regression models for hypothesis testing has been pointed out from a decade notably for seed germination (McNair et al., 2012; Onofri et al., 2019). Appropriate methods for time-to-event, also called *survival analysis* methods, widely developed in the medical field, start to be adapted to agriculture (Humplík et al., 2020; Romano and Stevanato, 2020). In this paper, we propose a phyllochron model in the spirit of these approaches, in which the phyllochron process is modelled as a succession of independent times-to-events corresponding to the times between appearance of successive leaves.

To evaluate the benefit of our approach, we make use of an original plant material that results from 12 years of two divergent selection experiments (DSE) for flowering time within two maize inbred lines under agronomical conditions (Durand et al., 2010, 2015). Within homogeneous genetic background, phenological shifts between Early and Late progenitors were selected, resulting in a difference reaching three weeks after 15 generations (150 degree-days, (Durand et al., 2015)). Representative progenitors from generation G13 were chosen to monitor plant growth during three years, between 2014 and 2016.

In addition to look for phyllochron differences between genotypes of the G13 generation, we investigated whether conditions experienced during growth could lead to alteration in the phyllochron. To address this question a modeling procedure allowing changes in leaf appearance rate was required. Indeed, although ATT is supposed to account for thermal dependency of growth rates, other climatic variables have been shown to modulate developmental rates in maize (Millet et al., 2019). Specifically to leaf appearance rates, a decrease of photosynthetically active radiation seems to slow down the phyllochron in a context of mixed cultivation systems (Zhu et al., 2014; Birch et al., 1998) while long photoperiod increases the leaf appearance rate (Warrington and Kanemasu, 1983; Miralles and Richards, 2000). Water and nitrogen stresses have also been shown to impact leaf elongation rate and to slow down the phyllochron (Chenu et al., 2008; Longnecker and Robson, 1994). To a lesser extent, sowing date is another factor affecting leaf appearance rate, which can be considered as the resulting effect of multiple climatic variables (Birch et al., 1998).

In this paper, we took advantage of the novel approach for phyllochron modelling to implement a procedure that accounts for experimental constraints, and allows an associated testing procedure. In addition, a two-step procedure was proposed to analyse the impact of climatic variables on departures from constant leaf appearance rate.

## 2 Materials and methods

### 2.1 Experimental design and data collection

#### 2.1.1 Plant Material

The plants used in these experiments were produced by 12 years of divergent selection for flowering time in two maize inbred lines MBS847 (MBS) and F252. In each inbred, two early and two or three late genotypes were formed by recurrently selecting and selfing the earliest and latest flowering plants in Early and Late selection populations respectively. In F252, early genotypes are named FE36 and FE39 and late ones FL27, FL317 and FL318. In MBS, early genotypes are named ME49 and ME52 and late ones ML40 and ML53. Details about of the selection and selfing processes are provided in Durand et al. (2010). In generation G13, contrasted flowering times were observed between Early and Late selection populations in both ancestral lines (Durand et al., 2015).

#### 2.1.2 Crop experiment

G13 plants from the nine genotypes were sown on Saclay’s Plateau (France) in 2014, 2015 and 2016. 25 seeds were sown in lines of 5.2m with distance between two lines of 0.8m. For each inbred line, the parcel was formed of several plots of 14 lines protected on both sides by a line of control plants. In 2014, three lines were sown per genotype. In 2015, three to five lines were sown per genotype. In both years, genotypes were randomly assigned to lines and plots. In 2016, a particular attention was brought to ME52 and ML40 and 12 lines of each were sown in two plots. To avoid seedlings predation from birds, experimental plots were protected with nettings installed immediately after sowing until most of the plants had at least 6 to 7 visible leaves. Then, from nettings removal to the emergence of panicule, the number of leaves of each plant were counted twice a week (every two to four days) corresponding to the time interval between the stages GRO:0007013 and GRO:0007015 of the Cereal Plant Development Ontology (https://bioportal.bioontology.org/ontologies/GRO-CPD). Tip emergence of a leaf was considered as the leaf appearance criterion. To avoid evaluation mistakes due to leaf degradation, ranks of the third leaf and every odd new leaf following nettings removal were marked on the leaf with a pencil (requiring a brief nettings removal for rank 3).

A portion of the observed plants were dissected for further analysis, around a median leaf rank of 8.5 (year 2014), 10 (year 2015) and 9 (year 2016). These partially observed plants were used to infer the phyllochron model but their contribution is limited to the estimation of the first leaves appearance.

Altogether, we collected data on 1795 plants. For each plant, the leaf rank of the youngest visible leaf was recorded at various time points. The appearance of the last two leaves was not modelled and the corresponding records were removed. Note that the time of appearance of each leaf, which characterizes the phyllochron, is not observed; the only information is a time interval in which each leaf appears. Moreover, plants with a decreasing number of leaves between two successive time points were deleted as presenting errors of measurement. In 2014, data on the full phyllochron were collected for 318 plants with 8-17 [mean 12.5] observation time points. In 2015, data on the full phyllochron were obtained for 371 plants with 8-21 [mean 13] time points, and partial measurements for 328 plants. In 2016, data on the full phyllochron were available for 196 plants with 6-15 [mean 11.2] time points, and partial measurements for 233 plants.

#### 2.1.3 Climate variables

Daily climate variables were extracted from records on a hourly basis at the climatic station hosted at field location, downloaded stored in the climatik INRAE database (https://intranet.inrae.fr/climatik - Gif-Sur-Yvette station - number 91272003). Climate variables are described in Table 1. Mean temperature of the day was used to compute thermal time with the procedure described in (Parent et al., 2010) with the parameters estimated in (Parent and Tardieu, 2012).

**Table 1:** Climate variables collected from the INRAE database Climatik. Gif-sur-Yvette station - number 91272003 (https://intranet.inrae.fr/climatik)

| Climate variable | Unit | Abbrev. |
| --- | --- | --- |
| Temperature range | °C | TR |
| Penman evapotranspiration | *mm* | PETP |
| Calculated global radiation | *J.cm<sup>-2</sup>* | CGR |
| Global radiation | *J.cm<sup>-2</sup>* | GR |
| Photosynthetically active radiation | *J.cm<sup>-2</sup>* | PAR |
| Calculated sunshine duration | *hours* | CID |
| Pourcentage of sunshine | % | IF |
| Rain falls | *mm* | RF |
| Maximum rain falls | *mm.h<sup>-1</sup>* | RFX |
| >40% humidity duration | *hours* | HD4 |
| Wetness duration | *hours* | WD |
| Maximum humidity | % | HX |
| >90% humidity duration | *hours* | HD9 |
| Minimum humidity | % | HN |
| >80% humidity duration | *hours* | HD8 |
| Atmospheric humidity | % | H |
| Wind speed | *m.s<sup>-1</sup>* | W |
| Maximum wind speed | *m.s<sup>-1</sup>* | WX |
| Dewpoint temperature | °C | DT |
| Mean vapor pressure | *mbar* | MVP |
| Actinothermic index 10cm | °C | AI1 |
| Actinothermic index 50cm | °C | AI5 |
| Minimum temperature | °C | TN |
| Mean temperature | °C | T |
| Calculated mean temperature | °C | CMT |
| Maximum temperature | °C | TX |
| Maximum actinothermic index 10cm | °C | I1X |
| Maximum actinothermic index 50cm | °C | I5X |

### 2.2 Statistical model for the phyllochron

The phyllochron process of a plant is characterised by the time intervals between appearance of successive leaves, or equivalently the times of leaf appearance. In the context of repeated spaced measurements, these quantities are unobserved (fig. 1A) and we define a statistical phyllochron model that, combined with an inference algorithm, enables us to estimate the average phyllochron at the genotype level.

We denote by *lsg* the *g^th^* genotype from selection population *s* (Early or Late) in ancestral line *l* (F252 or MBS). Phyllochron model was implemented separately for each year of experiment. All notations are summarised in Table 2.

**Table 2:** Summary of notations.

| Notation | Description |
| --- | --- |
| $y$ | Index for year |
| $p$ | Index for plant |
| $lsg$ | Index for the $g^{th}$ genotype in selection population $s$ from inbred line $l$ |
| $f$ | Index for leaf rank |
| $[f_{\min}^{y,lsg}, f_{\max}^{y,lsg}]$ | Largest interval of leaf ranks such that leaves $f_{\min}^{y,lsg} - 1$ and $f_{\max}^{y,lsg}$ are observed on at least 10 plants from genotype $lsg$ on year $y$. |
| $[f_{\min}^0, f_{\max}^0]$ | Common interval to all genotypes and years, equal to $\bigcap_{y,lsg} [f_{\min}^{y,lsg}, f_{\max}^{y,lsg}]$ |
| $F_{lsg}$ | Overall maximum number of leaves for genotype $lsg$ |
| $H_{y,lsg,p,f}$ | Time between sowing and appearance of leaf $f$ on plant $p$ from genotype $lsg$ on year $y$ (in degree-days) |
| $Y_{y,lsg,p,f}$ | Time between appearance of leaf $f - 1$ and $f$ on plant $p$ from genotype $lsg$ on year $y$ (in degree-days) |
| $\mu_{y,lsg,f}$ | Mean of $Y_{y,lsg,p,f}$ i.e. average interval between appearance of leaf $f - 1$ and $f$ for genotype $lsg$ on year $y$ (in degree-days) |
| $\mu_{y,lsg}^{C(f_{\min})}$ | Mean of $H_{y,lsg,p,f_{\min}}$ i.e. average time between sowing and appearance of leaf $f$ for genotype $lsg$ on year $y$ (in degree-days) |
| $\sigma_{y,lsg,f}$ | Standard deviation of $Y_{y,lsg,p,f}$ (in degree-days) |
| $\sigma_{y,lsg}^{C(f_{\min})}$ | Standard deviation of $H_{y,lsg,p,f_{\min}}$ (in degree-days) |
| $X_{y,c}(t)$ | Value of the climatic variable $c$ on day $t$ from year $y$ |
| $\bar{\mu}_{y,lsg,f}^{cal}$ | Average calendar time between sowing and appearance of leaf $f$ for genotype $lsg$ on year $y$ (in days) |

#### 2.2.1 Statistical model for the whole phyllochron process

Denote by *Y_y,lsg,p,f_* the unobserved time interval between appearance of leaves of rank (*f* – 1) and *f*, on plant *p* from genotype lsg and year *y*. The phyllochron process of plant *p* is characterised by the vector (*Y_y,lsg,p,f_*)_*f* = 1,…, *F^lsg^*_ with *F^lsg^* the overall maximum number of leaves for genotype *lsg*. Then, the time of appearance of leaf *f* on plant *p* from genotype *lsg* is

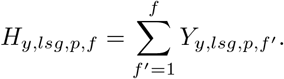

Notations are illustrated on Figure 1-A. We assume that the time interval between successive leaves *Y_y,lsg,p,f_* depends on some characteristics of the plant genotype *lsg* and on the leaf rank *f* = 1,…, *F^lsg^*:

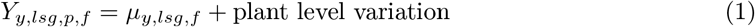

Thus, the average phyllochron of genotype *lsg* is characterized by the vector *μ_y,lsg_* = (*μ*_*y,lsg*,1_,… *μ_y,lsg,F^lsg^_*), and 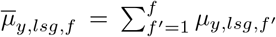 corresponds to the average time of appearance of leaf *f*. Note that no parametric form is assumed for the average phyllochron (*μ_y,lsg,f_*)*_f_*. Moreover, time intervals between appearance of successive leaves on each plant are assumed to be independent random variables and Gaussian distributed, thus

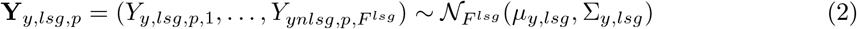

with 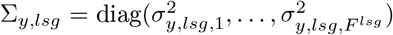 the diagonal variance-covariance matrix. The main parameters of interest are (*μ_y,lsg,f_*)*_f_*, while (*σ_y,lsg,f_*)*_f_* may be considered as nuisance parameters. Besides, note that the times (*H*_*y,lsg,p*,1_,…, *H_y,lsg,p,F^lsg^_*) of leaf appearance on a plant *p* are not independent, as they result from accumulation of independent intervals between leaves (*Y_y,lsg,p,f_*)*_f_*.

#### 2.2.2 Taking into account experimental constraints: restricted range of leaf ranks

Observations in the field started at a fixed time point, resulting in variations of the first leaf rank observed between plants. Moreover, a leaf rank has to be observed on a minimum number of plants in order to get correct estimates of the corresponding phyllochron parameters. Thus for each genotype lsg, we modelled the phyllochron on the range of leaf ranks 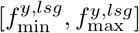 such that leaf ranks 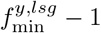 and 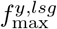 were observed on at least 10 plants from this genotype on year *y*.

Genotypes displayed differences in leaf ranks intervals which may originate both from statistical sampling and from biological variations. Nevertheless, some analyses required a common interval of leaf ranks, so we also considered 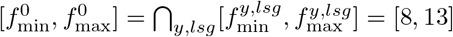. But this restricted interval leads to a loss of information. Therefore, comparative analyses between genotypes were performed using the common interval, while the temporal trends in phyllochron dynamics as well as the influence of climate were studied on the intervals 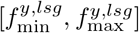.

#### 2.2.3 Statistical model on the restricted leaf rank interval: cumulated and instant phyllochron

First leaves being unobserved or with few observations, the corresponding parameters (*μ_y,lsg,f_, σ_y,lsg,f_*) cannot be estimated. Therefore, for a leaf rank interval [*f*_min_, *f*_max_], than can be equal to 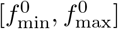 or 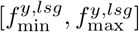 we can infer the following distributions (Figure 1-B).

- The distribution of 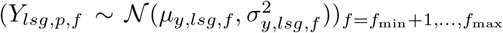, that we denote *instant phyllochron*.
- The distribution of 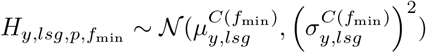 with

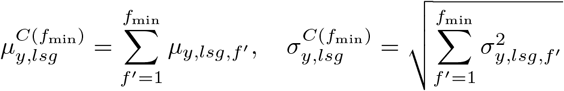

that we denote *cumulated phyllochron*. This quantity is the resultant of the plant development since sowing. Note that the distinction between cumulated and instant phyllochron is linked to the experimental conditions: if leaf number recording had started later in development, the cumulated phyllochron would be the resultant of more developmental stages.

#### 2.2.4 Parameter inference

For each genotype, the parameters

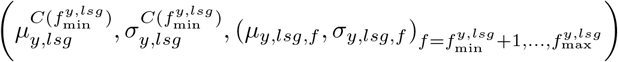

were estimated via a Monte Carlo Expectation Maximisation algorithm (Figure 2 and Suppl. Mat. A). Parameters of the cumulated phyllochron for the restricted interval 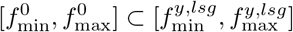 can be directly deduced. Indeed, if 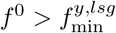:

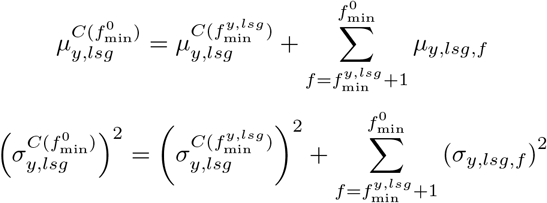

**Figure 2:**
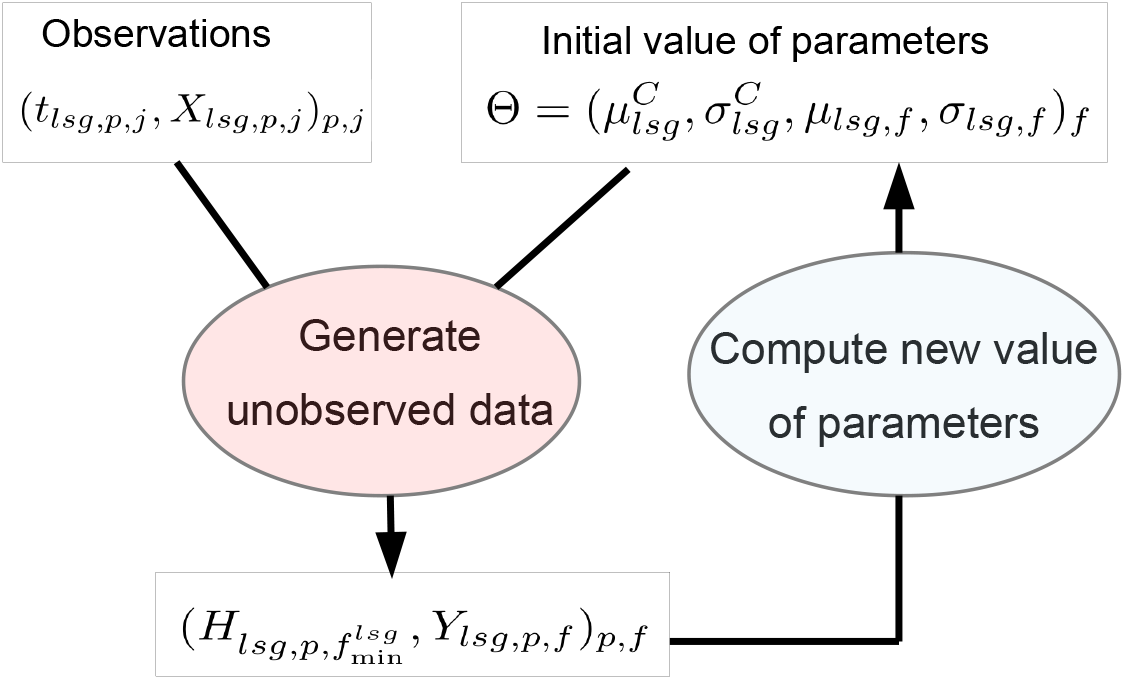
Monte Carlo Expectation Maximisation algorithm. Starting from an initial value of the parameters, the unobserved times of leaf appearance are drawn from their distribution given the observed data. Then, the maximum likelihood estimator is inferred from the simulated unobserved data, producing a new estimate of the parameters. The algorithm is iterated until stabilisation of the parameters.

### 2.3 Model comparisons

#### 2.3.1 Tests of genotypic groups effects

We made use of the hierarchical grouping of plants to understand the genetic factors that impact phyllochron. Indeed, differences between plants may come from ancestral lines (F252 versus MBS), selection population (Early versus Late), or genotypes within a selection population and an ancestral line. Note that because of the hierarchical structure of the models, significant differences between genotypes may result in significant differences between selection populations, independently of a divergent selection effect. In addition, statistical differences can occur either before the first observations and target the *μ^C^* parameter, or on the modeled leaf ranks and target the μ parameters. Altogether, on the leaf ranks common to all genotypes 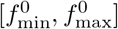, we ran seven different models *M_ij_* (Table 3), indexes *i* (resp. *j*) denoting the level of dependence of cumulated (resp. instant) phyllochron respectively. *i* (resp. *j*) values indicates that cumulated (resp. instant) phyllochron is considered as identical accross all plants (0), within the ancestral line (1), within the selection population (2) or within a genotype (3).

**Table 3:** Models for genotypic groups effects. C: cumulated phyllochron, I: instant phyllochron.

| Model | Cumulated phyllochron parameters | Instant phyllochron parameters | Model name |
| --- | --- | --- | --- |
| $M_{00}$ | $\forall(l, s, g), (\mu_{lsg}^C, \sigma_{lsg}^C) = (\mu^C, \sigma^C)$ | $\forall(l, s, g), (\mu_{lsg}, \sigma_{lsg}) = (\mu, \sigma)$ | identical |
| $M_{10}$ | $\forall(s, g), (\mu_{lsg}^C, \sigma_{lsg}^C) = (\mu_l^C, \sigma_l^C)$ | $\forall(l, s, g), (\mu_{lsg}, \sigma_{lsg}) = (\mu, \sigma)$ | C-line |
| $M_{11}$ | $\forall(s, g), (\mu_{lsg}^C, \sigma_{lsg}^C) = (\mu_l^C, \sigma_l^C)$ | $\forall(s, g), (\mu_{lsg}, \sigma_{lsg}) = (\mu_l, \sigma_l)$ | (C+I)-line |
| $M_{21}$ | $\forall g, (\mu_{lsg}^C, \sigma_{lsg}^C) = (\mu_{ls}^C, \sigma_{ls}^C)$ | $\forall(s, g), (\mu_{lsg}, \sigma_{lsg}) = (\mu_l, \sigma_l)$ | C-selection |
| $M_{22}$ | $\forall g, (\mu_{lsg}^C, \sigma_{lsg}^C) = (\mu_{ls}^C, \sigma_{ls}^C)$ | $\forall g, (\mu_{lsg}, \sigma_{lsg}) = (\mu_{ls}, \sigma_{ls})$ | (C+I)-selection |
| $M_{32}$ | $(\mu_{lsg}^C, \sigma_{lsg}^C) = (\mu_{lsg}^C, \sigma_{lsg}^C)$ | $\forall g, (\mu_{lsg}, \sigma_{lsg}) = (\mu_{ls}, \sigma_{ls})$ | C-genotype |
| $M_{33}$ | $(\mu_{lsg}^C, \sigma_{lsg}^C) = (\mu_{lsg}^C, \sigma_{lsg}^C)$ | $(\mu_{lsg}, \sigma_{lsg}) = (\mu_{lsg}, \sigma_{lsg})$ | (C+I)-genotype |

Model *M*_00_ assumes that all plants have the same phyllochron distribution whatever the ancestral line, the selection population or the genotype. Models *M*_10_ and *M*_11_ suppose that phyllochron varies with the ancestral line for at least one leaf rank, either during early developmental stages (*M*_10_) or anywhere throughout the season (*M*_11_); *M*_11_ is preferred to *M*_10_ when the phyllochron differences occur on observed leaf ranks. Similarly, models *M*_21_ and *M*_22_ assume differences in phyllochron between Early and Late populations for at least one leaf rank, while models *M*_32_ and *M*_33_ assume differences between genotypes.

As phyllochron estimates highlight obvious differences between years of experimentation, a separate analysis was run for each year. Also, comparisons between *M*_12_/*M*_22_ and *M*_11_ were performed either by pooling the two ancestral lines, or within each ancestral line. Similarly for comparisons between *M*_23_/*M*_33_ and *M*_22_ within ancestral line and selection population.

#### 2.3.2 Parametric sub-models of the phyllochron dynamics

In order to elucidate trends in the departure from the classic constant leaf rate appearance model, we considered parametric models for the instant phyllochron as a function of leaf rank, that can capture general trends by summarizing the vector 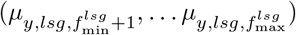 through a reduced set of parameters for each genotype and year. We considered four parametric models. (i) The constant model supposes that phyllochron is constant throughout the season. (ii) The constant-rate model supposes a constant increase or decrease of phyllochron with leaf rank. (iii) The piecewise constant model allows a single change in the phyllochron at a given leaf stage. (iv) The piecewise linear model allows two increasing or decreasing phases. Details on model selection are provided in Suppl. Mat. B.

#### 2.3.3 Criteria for model comparison

We considered the *χ*^2^-likelihood ratio test, as well as the AIC and BIC criteria (Suppl. Mat. D). As these criteria rely on asymptotic considerations and may be biased in the context of a finite sample size, we validated them by a complementary permutation test. The instant phyllochrons of genotypes FE36 and FE39 on year 2015 were compared, with the null assumption of equal phyllochron distribution for the two genotypes. Then the same comparison was repeatedly performed while permutating the genotype labels and thus defining “false genotypes”; if the likelihood ratio test was unbiased, the p-value for the “false genotypes” would be uniformly distributed on [0, 1]; moreover, the most relevant criterion among AIC and BIC should mostly select the model under the null assumption.

#### 2.3.4 Principal Components Analysis (PCA)

PCA was implemented on the set of vectors

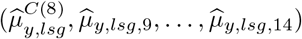

for all combinations of genotypes *lsg* and all years *y*. For genotypes FE36 and FL317 in 2014, 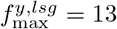 so 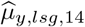 was not estimated, and we replaced these two missing values by the average of 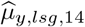 over all other genotype-year combinations.

### 2.4 Impact of climate on phyllochron

In order to evaluate the impact of climate on the variations of phyllochron throughout the season and between years, we considered a model where the interval times between successive leaves were regressed over a function of longitudinal climatic variables prior to the leaf appearance.

#### 2.4.1 General model

Our climate model is performed on the calendar time scale. Therefore, phyllochron parameters are first back-transformed into their calendar time counter part. Thus 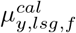 and 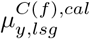 denote respectively the average time between appearance of leaves (*f*–1) and *f* and the average appearance time of leaf *f* since sowing on genotype *lsg*, on calendar time. Then, for year *y*, genotype *lsg* and leaf rank *f*, we consider the general model:

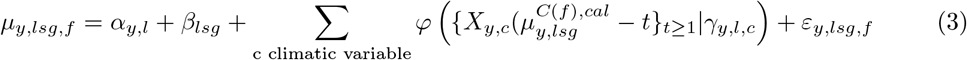

where 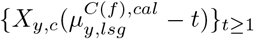 is the vector of values of the climatic variable *c* on year y prior to the average appearance time of leaf *f* on genotype *lsg*, and *φ*(·|*γ*) is a parametric function which characterises the impact of the longitudinal climatic variables on the phyllochron (Figure S3-A).

The line-year effect *α_y,l_* accounts for the potential effect of genetic background interacting with the year of experiment (climatic and cultivation conditions and sowing date). The effect of each climatic variable 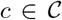 depends on the year-line background through the coefficient *β_y,l,j_*. Moreover, the effect of genotype is accounted for through the coefficient *γ_lsc_* identical over years, and residuals *ε_y,lsg,f_* are assumed independent identically distributed (i.i.d.) centered Gaussian.

#### 2.4.2 Parameterization

The function *φ* was parameterized as piecewise constant:

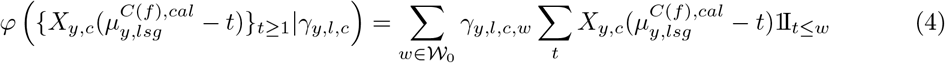

This flexible parameterization can model various phenomena with a small number of non-zero coefficients (*γ_y,l,c,w_*), depending on the coefficient values (Figure S3-B). We considered the set of time windows 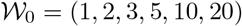 days, to allow both a fine modelling of short time effects and potential long term impacts of the climatic variables.

#### 2.4.3 Inference with lasso regression

The model was inferred by replacing unknown true phyllochron parameters (notations “*μ*”) by their estimates computed by the Monte-Carlo EM algorithm (notations 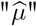). To account for the large number of parameters in model (3) with parameterization (4), inference was performed by a lasso regression, which automatically sets to zero a large proportion of the coefficients by means of a *L*^1^-penalty; this proportion is tuned via a constant in the penalty. We used the R package `penalized` which allows to select the optimal penalty constant based on cross-validation. In order to preserve the data structure, cross-validation splitting was implemented by keeping together phyllochron measures of each genotype-year combination.

The model performances were assessed through the prediction error. More precisely, for each year-genotype combination (*y, lsg*), the climatic model (3, 4) was inferred using the measurements of plants from all genotypes-years (*y′, lsg′*) ≠ (*y, lsg*); then the predicted values of the phyllochron parameters 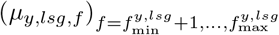 were computed based on (3); finally the Mean Squared Error (MSE) was computed between these predicted values and the values inferred in section 2.2, considered as the “true” values.

#### 2.4.4 Permutation-like test

The MSE described in the previous sub-section may over-estimate the model prediction ability since the same climate was used in training and validation sets. To correct this bias, MSE was repeatedly computed using “false” climates whose seasonal variations should not be associated with phyllochron variations. False climate were generated using climate records on years 2009 to 2017 on three periods of time, leading to 252 false climates. Then, the proportion of “false” climate for which the MSE is smaller than the one obtains with the “true” climate stands for a p-value of the impact of climate on phyllochron. Indeed, a weak proportion indicates that the climate partly explains the phyllochron variations out of potential statistical bias.

## 3 Results

### 3.1 Total leaf number

Variations in total leaf number were observed (Figure S1). Early genotypes have globally less leaves than Late genotypes from the same genetic background, and this difference is stronger for the ancestral line MBS with 16-18 leaves versus 18-20 leaves. Besides, the total leaf number is similar across years within genotypes, even slightly higher in 2014 than in 2015, and in 2015 than in 2016 except for genotype FL317. By construction, the maximum modelled leaf rank 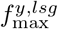 is correlated to the total leaf number, thus the trends observed on the latter are recovered on the former.

### 3.2 Phyllochron estimates by genotypes

The data structure enables to conduct a hierarchical analysis of each grouping level effect: year, ancestral line, selection population and genotype.

#### 3.2.1 Descriptive analysis of cumulative phyllochron and global instant phyllochron

Figure 3 displays the estimates of the cumulated phyllochron between sowing and the appearance of the 8*^th^* leaf as well as the interval of time between leaves 8 and 13 in 2014 and 2015.

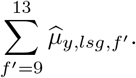

We observe that both cumulated and instant phyllochrons are strongly impacted by year, but in opposite sense: While the cumulated phyllochron (Figure 3A) is faster in 2015 than in 2014 (difference of 3-5 degree-days), the global instant phyllochron (Figure 3B) is faster in 2014 and slower in 2015 (difference of 2-3 degree-days); Besides, on the instant phyllochron, differences between years are more marked for ancestral line MBS (difference of 3-7 degree-days) than F252 (1-2 degree-days).

**Figure 3:**
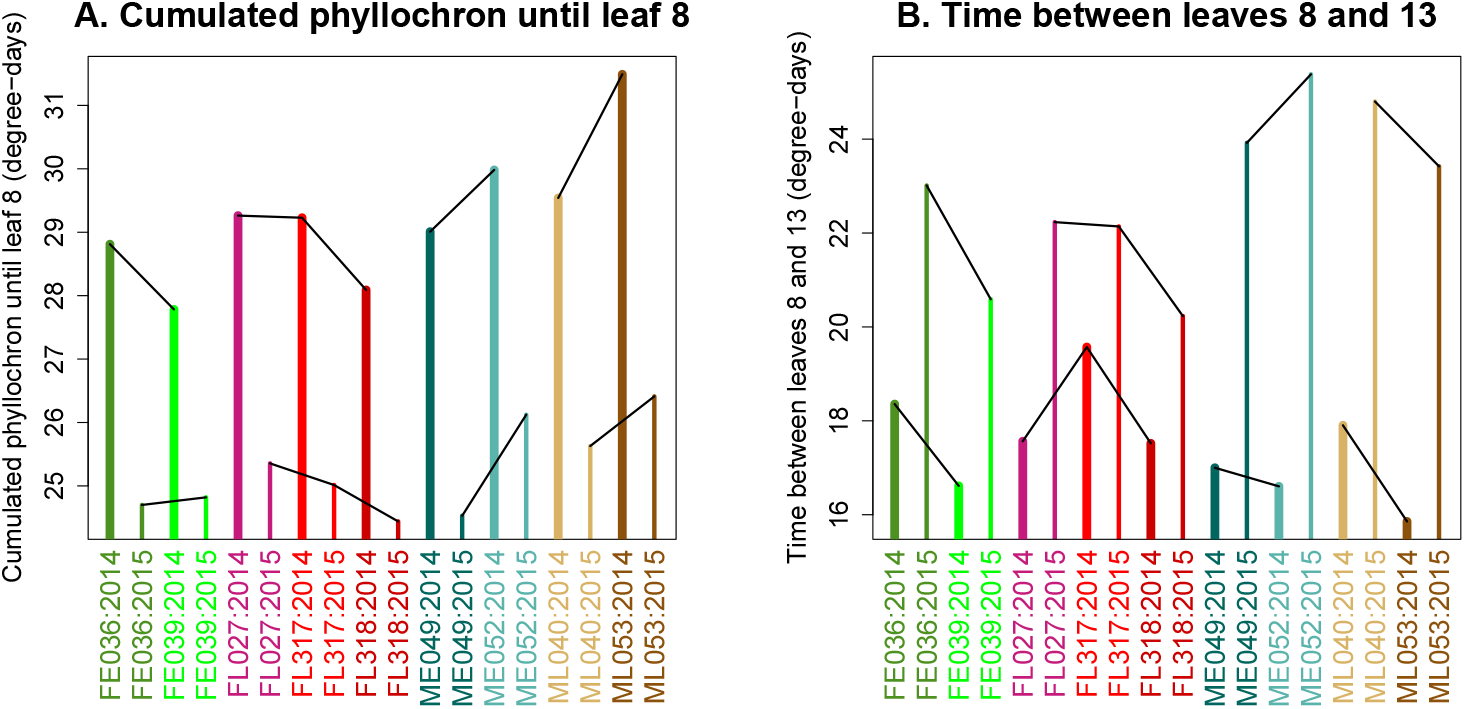
Cumulated and instant phyllochron. **A.** Accumulated thermal time (ATT) in degree-days to reach eight visible leaves. **B.** Time interval in degree-days between appearance of leaves eight and 13. Straight lines join genotypes from the same selection population observed the same year. Thin bars corresponds to year 2015 and thick bars to 2014.

Regarding ancestral line effect, the cumulated phyllochron is faster for F252 (28.7 and 24.8 degree-days in 2014 and 2015) than for MBS (30.2 and 25.7 degree-days in 2014 and 2015) for both years. This means that genotypes from the F252 background reach eight visible leaves earlier than genotypes from the MBS background. Concerning the global instant phyllochron, <u>i.e</u> the ATT between leaves 8 and 13, differences between ancestral lines depend on the year of observation. In 2015, it took more time for MBS genotypes (23.7 degree-days) than for F252 genotypes (21.5 degree-days). The reverse was observed in 2014: F252 genotypes took 17.8 degree-days between the appearance of leaves eight and 13, while MBS genotypes took 16.7 degree-days.

No general trend is visible between selection populations. However, we observe differences between genotypes from the same selection population. Interestingly, those differences tend to be preserved from one year to the next: FE039 tends to be faster than FE036 (except for cumulated phyllochron in 2015), FL318 tends to be faster than FL317, and ME049 tends to be faster than ME052 (except for instant phyllochon in 2014). The two mentioned exceptions correspond to weak absolute value of the difference: 0.11 degree-days between cumulated phyllochron of FE36 and FE39 in 2015 and 0.39 degree-days between instant phyllochron of ME49 and ME52 in 2014, which makes the change of trend between 2014 and 2015 poorly informative. Concerning the Late MBS genotypes, we observe a difference between instant and cumulated phyllochron: ML040 tends to be faster than ML053 until eight leaves, and longer between 8 and 13 leaves.

Figure 4 provides a more detailed insight on the instant phyllochron dynamics, by showing the estimations of the instant phyllochron for all leaf ranks for which data were available in the different genotypes. Again, the most stricking observation is the strong variation of the temporal trend between years. In 2014 and 2016, time interval between successive leaves varies moderately between 3 and 5 degree-days, except an increasing trend between leaves 13 and 14 in F252 genotypes, and is globally shorter in 2014 than in 2016 (3.5 versus 4 degree-days on average). In 2015, the phyllochron shows stronger variations throughout the season, between 3 and 7 degree-days, with an increasing and a decreasing phase for MBS genotypes, and a rather constant increase for F252 genotypes. Note that the specific temporal pattern observed in 2015 in Figure 4 explains the high values of the ATT between leaves 8 and 13 observed in Figure 3.

**Figure 4:**
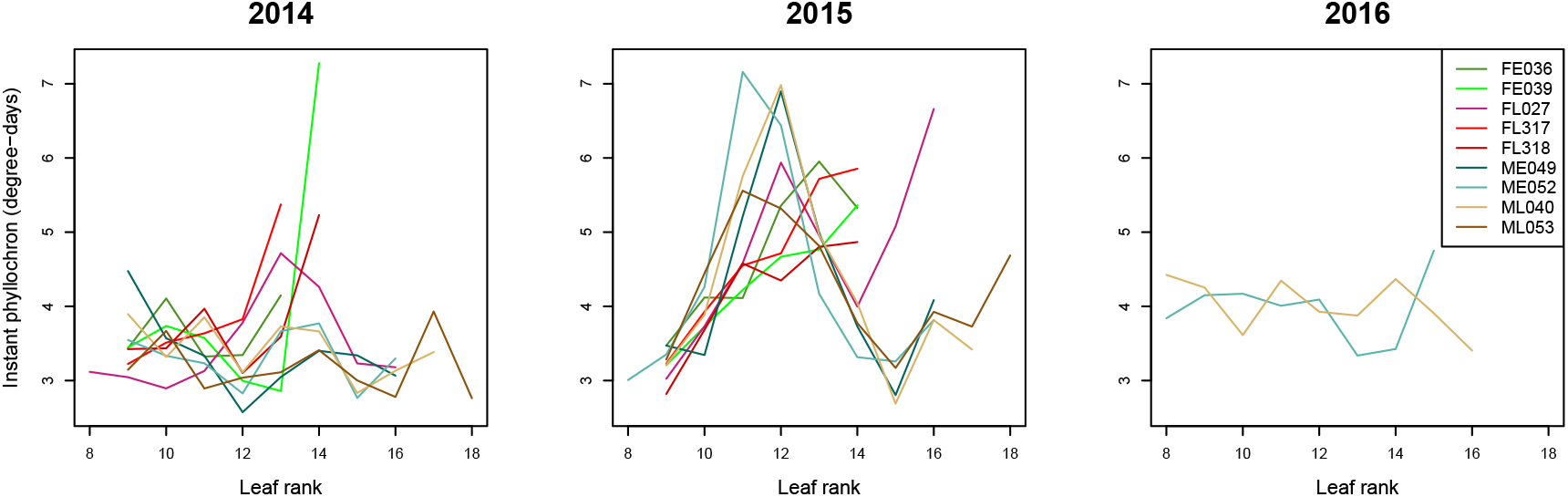
Estimates of instant phyllochron. Each plot corresponds to a year; Colors correspond to the different genotypes; Lines join estimates 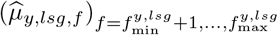 from the same genotype *lsg* at successive leaf ranks.

#### 3.2.2 Model comparisons

Genotypic groups comparisons were performed only for years 2014 and 2015, where all genotypes were observed, and were restricted to the range of leaf ranks common to all genotypes. The preliminary permutation test analysis indicates that the *χ*^2^-test and the AIC criterion are reliable for group effect testing, even if AIC is slightly biased in favor of the largest model (Suppl. Mat. D).

Table 4 displays the result of comparisons between ancestral lines, selection populations and genotypes. Both criteria mostly provide coherent conclusions, while AIC is slightly less conservative as highlighted by the permutation test.

**Table 4:** AIC criterion, and *χ*^2^-likelihood ratio p-value for ancestral line, selection and genotypic effects. For each grouping effect, models *M*_*j*–1,*j*–1_, *M*_*j*–1,*j*_, and *M_j,j_* are compared (cf Table 3). Bold numbers correspond to smallest AIC value and *χ*^2^-likelihood ratio *p* < 0.01. Best model consists in either no group effect (ns), a significant group effect on cumulative phyllochron (C), a significant group effect on instant phyllochron (I) or both (C+I)

Ancestral line effect
| | AIC-$M_{j,j}$ | AIC-$M_{j,j-1}$ | AIC-$M_{j-1,j-1}$ | $\chi^2-M_{j,j}/M_{j-1,j-1}$ | $\chi^2-M_{j,j}/M_{j,j-1}$ | $\chi^2-M_{j,j-1}/M_{j-1,j-1}$ | best |
| --- | --- | --- | --- | --- | --- | --- | --- |
| 2014 | **3544** | 3605 | 3649 | **<e-16** | **3.20e-13** | **1.70e-11** | C+I |
| 2015 | **7453** | 7662 | 7709 | **<e-16** | **<e-16** | **2.60e-12** | C+I |

Selection effect
| | AIC-$M_{j,j}$ | AIC-$M_{j,j-1}$ | AIC-$M_{j-1,j-1}$ | $\chi^2-M_{j,j}/M_{j-1,j-1}$ | $\chi^2-M_{j,j}/M_{j,j-1}$ | $\chi^2-M_{j,j-1}/M_{j-1,j-1}$ | best |
| --- | --- | --- | --- | --- | --- | --- | --- |
| 2014 | **3489** | 3542 | 3544 | **8.10e-12** | **2.00e-11** | 4.60e-02 | I |
| 2014.F | **2182** | 2238 | 2239 | **2.00e-12** | **2.60e-12** | 8.60e-02 | I |
| 2014.M | 1307 | **1303** | 1304 | 4.20e-02 | 7.20e-02 | 7.20e-02 | ns |
| 2015 | **7427** | 7441 | 7453 | **4.90e-07** | **4.30e-05** | **3.20e-04** | C+I |
| 2015.F | **4252** | 4260 | 4258 | **3.00e-03** | **1.70e-03** | 9.00e-01 | I |
| 2015.M | **3175** | 3181 | 3195 | **1.30e-05** | **2.90e-03** | **6.00e-05** | C+I |

| | AIC-$M_{j,j}$ | AIC-$M_{j,j-1}$ | AIC-$M_{j-1,j-1}$ | $\chi^2-M_{j,j}/M_{j-1,j-1}$ | $\chi^2-M_{j,j}/M_{j,j-1}$ | $\chi^2-M_{j,j-1}/M_{j-1,j-1}$ | best |
| --- | --- | --- | --- | --- | --- | --- | --- |
| 2014 | **3402** | 3462 | 3489 | **<e-16** | **1.0e-13** | **8.2e-07** | C+I |
| 2014.Fearly | **758** | 767 | 774 | **6.5e-05** | **1.1e-03** | **2.5e-03** | C+I |
| 2014.Flate | **1352** | 1399 | 1408 | **5.6e-12** | **2.1e-10** | **1.5e-03** | C+I |
| 2014.Mearly | **681** | 677 | 677 | 6.0e-02 | 7.6e-02 | 1.4e-01 | ns |
| 2014.Mlate | **612** | 619 | 629 | **3.8e-05** | **1.8e-03** | **5.3e-04** | C+I |
| 2015 | 7266 | **7353** | 7427 | **<e-16** | **<e-16** | **1.1e-16** | C+I |
| 2015.Fearly | **1904** | 1917 | 1915 | **4.9e-04** | **2.6e-04** | 9.4e-01 | I |
| 2015.Flate | **2297** | 2323 | 2337 | **3.7e-09** | **7.3e-07** | **1.4e-04** | C+I |
| 2015.Mearly | **1948** | 1994 | 2044 | **<e-16** | **2.6e-10** | **4.9e-13** | C+I |
| 2015.Mlate | **1117** | 1119 | 1131 | **1.5e-04** | **1.0e-02** | **2.6e-04** | C+I |

The results indicate a strong effect of the ancestral line on both cumulated and instant phyllochron and confirm the previous graphical observations of Figure 3: genotypes derived from F252 or MBS behave differently for the phyllochron. Difference between selection populations are more difficult to interpret because they depend on year and ancestral line. In 2014, we found significant differences for instant phyllochron between Early and Late populations of the F252 ancestral line, that may explain the significance of the general comparison. No significant difference was found between Early and Late MBS populations. In 2015, we found significant differences for instant phyllochron between Early and Late populations in all ancestral lines. We also found a significant difference between Early and Late cumulated phyllochron in the MBS ancestral line, that may explain the significance of the general comparison. Finally, almost all comparisons between genotypes within selection lines are significant. The only exceptions were the Early MBS genotypes that behave similarly in 2014, and the Early F252 genotypes that exhibit the same cumulated phyllochron in 2015. Those results confirm the graphical observations in Figure 3. They show that, concerning the phyllochron, each genotype can be characterized by a specific temporal pattern and that the statistical model was able to retrieve significant differences.

#### 3.2.3 Graphical analysis of instant phyllochron

Significant differences with statistical tests can be interpreted by a graphical analysis of the estimates of instant phyllochron (Figure 4). Differences between ancestral lines are mostly visible in year 2015: ATT between successive leaves increases with leaf rank in F252, while it shows a maximum around 11-12 leaves in MBS. In 2014, the tendency to increase is conserved in F252 genotypes, while MBS genotypes seem to be constant or decreasing. Regarding differences between selection populations, the most visible effect concerns the difference between Early and Late F252 genotypes, with a constant instant phyllochron in Early genotypes and an increasing phyllochron in late genotypes. Other differences affect particular leaf ranks. For example, instant phyllochron is higher at leaf rank 11 in Late F252 than in Early F252 genotypes in 2015, or at leaf rank 13 in Late MBS than in Early MBS genotypes in 2015.

Figure 5 focuses on the average temporal trends between leaves 9 and 13 for each genotype and each year according to the selection population. A single line is represented for Early MBS genotypes in 2014 because no significant difference was detected between the two genotypes. The statistical tests from section 3.2.2 indicate a genotype effect among most selection populations, but they do not tell about the structure of this effect. Figure 5 enables to refine this analysis and indicates that differences of phyllochron between genotypes do not have the same magnitude on all leaf ranks and may mostly be associated by a limited number of leaf ranks. For example, on the selection population ML2015, difference between the two genotypes is mostly observed on leaf 12. Other type of differences can be observed, such as for the Early population ME2015, for which phyllochron displays a lag between the two genotypes (the maximum phyllochron value being observed at leaf rank 11 for ME052 and at leaf rank 12 for ME049).

**Figure 5:**
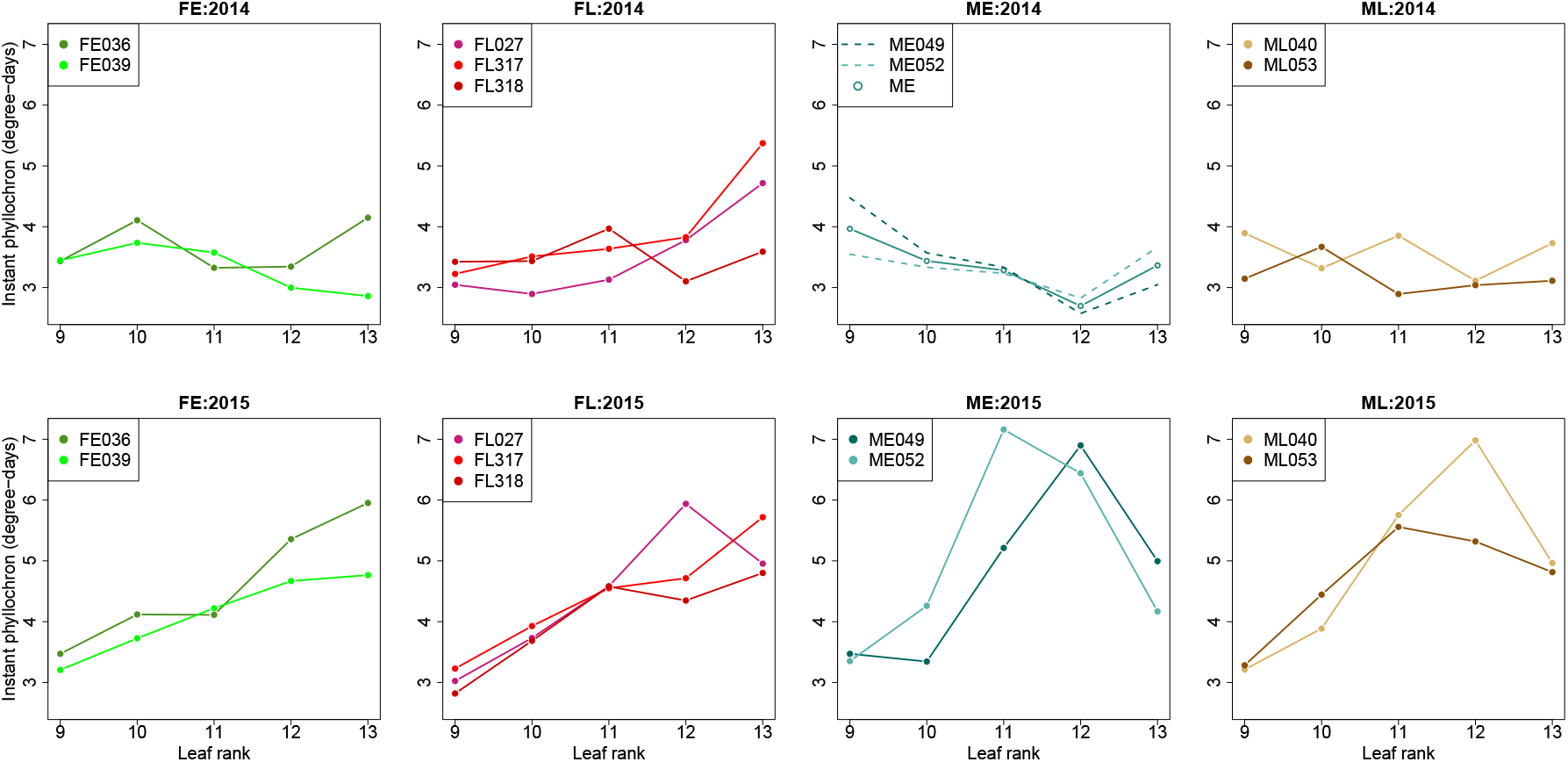
Genotypic and selection effects on instant phyllochron. Instant phyllochron of all genotypes within selection groups. Column corresponds to selection population and row to year. When differences between genotypes within selection population is not significant, the phyllochron estimate on the whole selection groups is represented.

#### 3.2.4 Summary of genotypic group effect analysis

Altogether, we found significant differences on phyllochron at all levels of grouping, for both cumulated and instant phyllochrons. While the year effect is predominant, significant differences were observed between ancestral lines on both cumulated and instant phyllochron, with MBS exhibiting an higher phyllochron, corresponding to a slower leaf appearance rate. Differences at lower levels (selection populations and genotypes) are also globally statistically significant but selection population effects may be driven by genotype effects, which seems confirmed by the graphical analyses which does not highlight selection population trends.

Moreover, we observe strong genotype-by-year interactions as differences of instant phyllochron between genotypes exhibit different patterns from one year to the other. Roughly, three different temporal patterns were observed: (i) constant phyllochron (Early F252 and all MBS genotypes in 2014), (ii) increasing phyllochron (Late F252 in 2014 and all F252 genotypes in 2015), (iii) increasing/decreasing phyllochron (all MBS genotypes in 2015). Those results motivated the test of parametric models.

### 3.3 Parametric models

The procedure to select the best parametric models ensures that the temporal trends observed on the instant phyllochron are significant and do not originate from sampling variations. Figure 6 displays the best parametric sub-model for each genotype. Overall, the sub-models fit well the complete models, and manage to capture the seasonal tendencies. The constant phyllochron model was selected only for one genotype in 2014 (ME052). Moreover, the major differences previously observed are recovered by the best parametric sub-model. In 2014 and 2016, the variations of phyllochron are moderate with a few exceptions in 2014 for the Late F252 genotypes, that increases either linearly (FL317, FL318) or piecewise (FL027). In 2015, the phyllochron of most F252 genotypes increases throughout the season, while the phyllochron of MBS genotypes and FL027 shows a maximum between 11 and 13 leaves, with an increase before and a decrease after.

**Figure 6:**
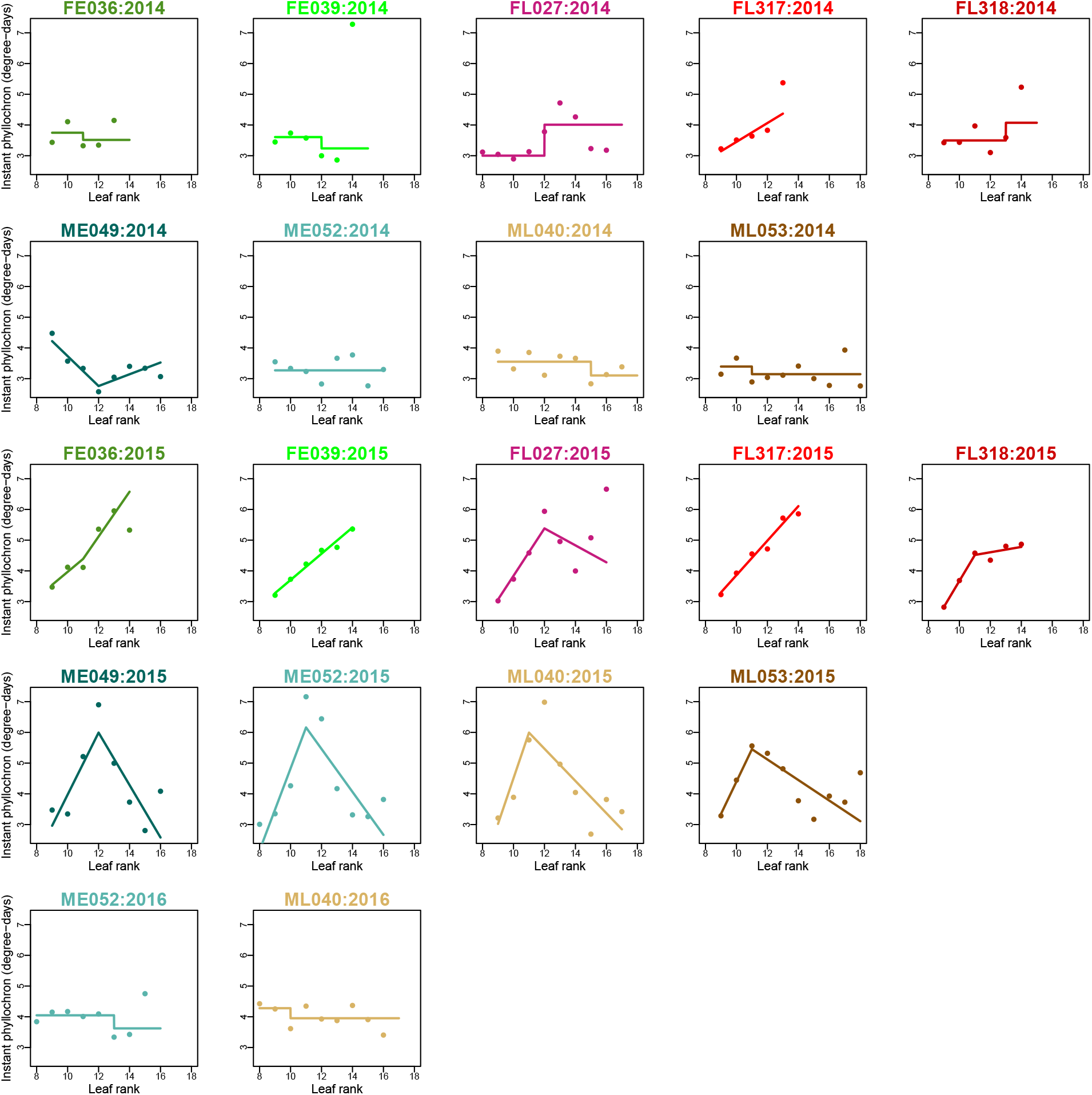
Best parametric sub-models. Each plot shows the instant phyllochron 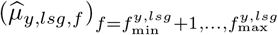 in degree-days for each genotype and each year. Dots correspond to the estimates under the complete model, while straight line displays the estimate under the best parametric sub-model.

### 3.4 Patterns of variation

PCA of all phyllochron estimates 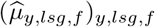 (Figure 7) provides an overview of global patterns. The first axis (49% of total inertia), discriminates year 2015 on the right versus years 2014 and 2016 on the left. The second axis (21% of total inertia) discriminates ancestral lines: F252 genotypes mostly have negative coordinates, while MBS genotypes have positive coordinates. No clear discrimination is observed between Early and Late genotypes within each ancestral line. Other differences concern genotypes, independently of the year of observation, the ancestral line or the selection effect. Besides, the cumulated phyllochron as well as the instant phyllochron of leaves 9 to 13 are loaded by both axes, and thus are associated with both ancestral line and year. On the contrary, 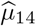 is mostly associated with ancestral line. Finally, phyllochron parameters are ordered clockwise by leaf rank in the correlation circle, which indicates that intervals between successive leaves are similar for close leaf ranks.

**Figure 7:**
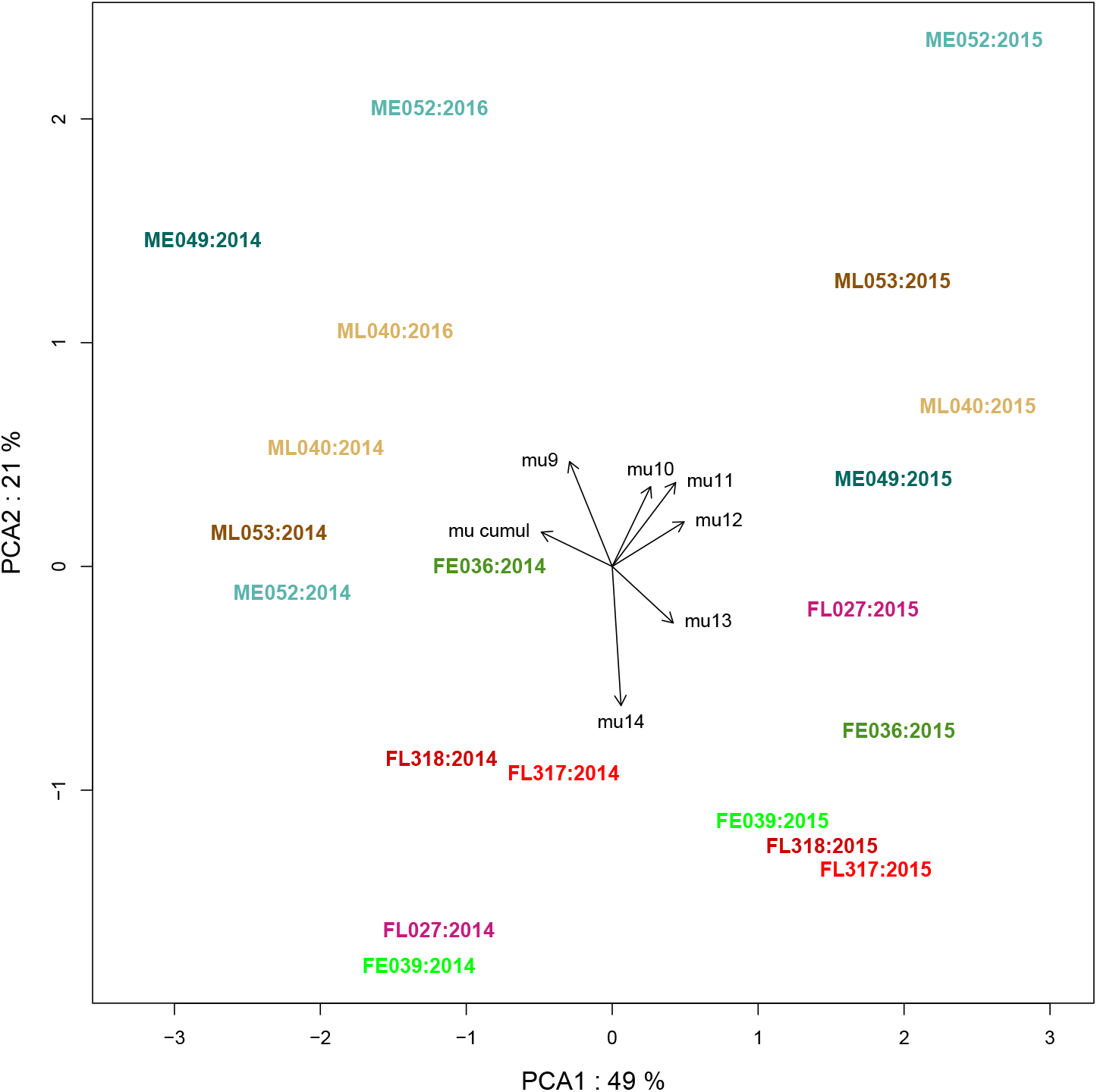
Principal Component Analysis of seasonal variations of the phyllochron. Variables are the average time 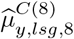 between sowing and appearance of leaf eight and the instant phyllochrons 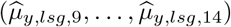 between leaf ranks nine and 14. Individuals are the genotype-year combinations. Individuals are represented on the first two axes of PCA and the colors correspond to the genotypes. Correlations between the variables and the PCA axes are represented by the black arrows using an arbitrary scale.

### 3.5 Input from climatic variables

Figure 8 displays the predicted values of phyllochron with the climatic model (3, 4) in a crossvalidation framework. The first row provides the phyllochron estimates in the three years and is similar to Figure 4. The second row gives the predictions from the climatic model. Whenever temporal trends are captured by predictions, this means that seasonal climate fluctuations are able to explain phyllochron variations. The third row shows the temporal fluctuations of the residuals of the climatic model. According to the climatic model, residuals are supposed to be independent of predicted values and should not exhibit any temporal pattern. Thus, Figure 8 indicates that significant temporal trends observed in 2015 are partially recovered by the climatic model. This is confirmed by the range of phyllochron variations in the residuals (−1/+1.5 degree-days) which is much smaller than the range of variation in the phyllochron estimates (3-7 degree-days). Nevertheless, the weaker variations are not captured by the model; for example, in 2016 the predicted phyllochrons are almost constant and all the observed temporal variation can be found in the temporal patterns of the residuals.

**Figure 8:**
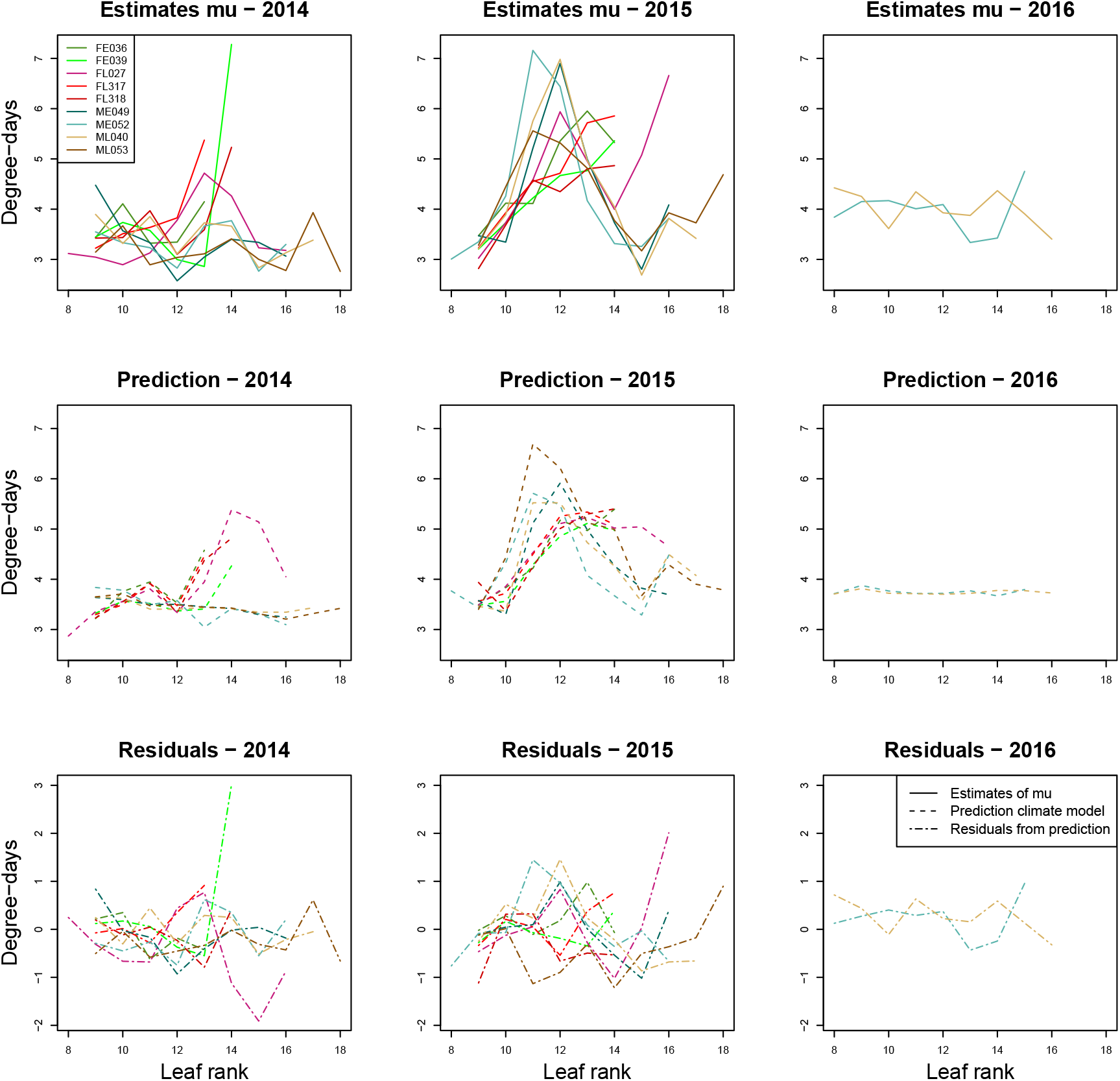
Predicted values and residuals of the climate model. Each plot corresponds to a year and each color to a genotype. The estimates (*μ_y,lsg,f_*) are recalled for comparison purpose on the first row; the second row displays the predicted values of 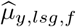 with the climate model (3 and 4) in a cross-validation framework, as a function of the leaf rank. The third row provides the residuals of prediction i.e. the difference between the two curves above.

As highlighted in Section 2.4.4, despite the cross-validation framework, the prediction of Figure 8 are not totally independent since the same climate records are involved in the training and the validation sets. Therefore, cross-validation predictions were computed with 252 “false” climates (see Section 2.4.4), and the Mean Square Error (MSE) defined as:

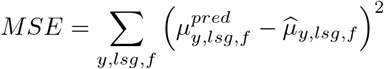

with 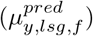 the phyllochrons predicted with the climatic model (3, 4) transformed in thermal time. The smaller the MSE, the better. The MSE was smaller with the true climate than with 94.8% of the false ones (Figure S4), which assessed a moderate predictive ability of the climatic model (*p* = 0.052).

Lasso regression may sometimes be used as feature selection by keeping variables with non-zero coefficients. However, in our context, this selection is not robust as the climatic variables are strongly correlated (Figure S5). Thus, we chose not to display a list that could lead to misleading interpretations. Instead, since phyllochron exhibits a peculiar structure in 2015, we decided to focus on this year in order to see if we could identify special patterns in climatic variables that might be hypothesized to explain the observations. Three variables, rainfalls, humidity and photosynthetic radiations attracted the attention (Figure S6) and year 2015 was characterized by low rainfall all along the season and two episodes of low humidity and high radiation: the first two weeks of June 2015 and the first week of July 2015. Therefore, 2015 was characterized by the establishment of progressive drought conditions for several weeks that might have contributed to the increase followed by a decrease in phyllochron this year. The specific phyllochron pattern observed in 2015 (and in last leaf ranks of a few genotypes in 2014) could originate from dry conditions.

## 4 Discussion

### 4.1 A stochastic process model for maize phyllochron

In this paper, we proposed a hypothesis testing model for the phyllochron based on a stochastic process in which phyllochron depends on leaf rank non-parametrically. This approach represents a more accurate modelling than the classic regression-based phyllochron models. Indeed, our results demonstrate strong variations of phyllochron throughout seasons, indicating that the linear phyllochron model assuming constancy, which is the most widely used model for hypothesis testing (Padilla and Otegui, 2005) may be too simple in the context of peculiar climatic conditions. Moreover, our analysis of various parametric sub-models suggests that a two-phase model could represent a reasonable compromise between simplicity and flexibility, even if the complete model is statistically more accurate. Bilinear models have been proposed in literature, for phyllochron analysis (Clerget and Bueno, 2013) or in prediction models (e.g. APSIM, Archontoulis et al. (2014)), but they do not enable hypothesis testing. Beyond modelling accuracy, our model allows to circumvent the statistical flaws of testing based on linear regression phyllochron.

Indeed, we proposed an unbiased statistical procedure to compare phyllochron in various conditions, that we applied to genotypic groups comparison. Generally, a price to pay for flexibility is the larger number of parameters, which requires more data to be correctly estimated, and this limitation is particularly true in the less informative context of interval censoring, in which information is restricted to time intervals in which leaves appear. Nevertheless, our results indicate that aproximately 40 plants by genotypes (year 2014) were sufficient to highlight significant differences between genotypic groups resulting from a recent divergence. However, in its actual version, our model cannot handle testing framework more complex than comparisons between classes or conditions.

The impact of climate on phyllochron variations through season was assessed with a two-step procedure: first, the parameters of the phyllochron model were inferred without accounting for climate, then climate variables cumulated before the appearance of each leaf were regressed on the average interval between leaves. Two-step inference has already been pointed as potentially biased in other statistical context (see e.g. Celeux (1989) in clustering). Notably the inference of the climatic model does not account for precision in the estimation of phyllochron parameters, but the use of a permutation-like test enables to limit the potential bias. Moreover, two-step inference remains a classic approximation when full-likelihood minimisation is not available. Notably, it is classically used in genotype-to-phenotype studies in which a dynamic phenotype is summarised by a restricted set of parameters on which statistical analyses are performed (Reymond et al., 2004; Marchadier et al., 2019).

Our model assumes Gaussian plant level variations around the genotype average phyllochron, which enabled us to use existing tools to sample from truncated multivariate distributions. Approximation of a positive variable by a Gaussian distribution is acceptable if the standard deviation is smaller than three-four times the mean, which is the case for most but not all genotypes on our data set. Nevertheless, Gaussian approximation is very common in statistical modelling, and it can be shown in diverse contexts that it leads to unbiased or moderately biased results.

This work is in the continuity of recent approaches for time-to-event analysis in agriculture (Humplík et al., 2020; Romano and Stevanato, 2020), as an alternative to flawed regression model but phyllochron modelling that include successive events is a much more complex framework. We are currently working on a more general framework based on semi-Markov models that that would directly include longitudinal covariates such as climatic variables within the phyllochron model. These models can also handle more general distributions and thus relax the normality assumption, and our first results indicates that the assumption of gaussian variations practically does not impact the average phyllochron estimates. Semi-markov models is a growing topic with diverse application domains, and future statistical and algorithm works could allow to include more complex covariate, and then handle, in an accurate modelling, testing frameworks that are nowadays addressed by constant phyllochron model.

### 4.2 Differences in phyllochron between closely related genotypes

Due to experimental constraints, we modelled phyllochron on a restricted range of leaf ranks (8-13 for global comparisons). Additionnally, the cumulated time between sowing and leaf 8 appearance constitutes an indicator of the first growing phase of the plant development, which cumulates emergence and the first phase of phyllochron.

As previously published, maize lines produced by Saclay’s DSE experiment exhibit a gradual flowering time divergence over the first 13 generations (Durand et al., 2010, 2015). The characterization of the phyllochron of these genotypes, as performed here, enables to better understand the developmental changes that could underlie such a response to selection. Our analyses does not highlight differences of phyllochron between selection populations (out of genotype-level effect). Nevertheless, we observed that the total leaf number was impacted during the selection process, with Late genotypes tending to produce more leaves than Early genotypes and a change in leaf number would be sufficient to accelerate or delay the flowering time (Durand et al., 2012). Besides, the small number of genotypes per selection population may be a limitation to distinguish confounded genotype and selection population effects.

A variability of phyllochron has already been reported between genetically distant maize inbred lines (Verheul et al., 1996) or between hybrids (Padilla and Otegui, 2005). Interestingly, our results demonstrate phyllochron differences between closely related genotypes and notably between genotypes selected in response to the same selective pressure (earliness or lateness), suggesting that the phyllochron evolved independently of the selection pressure for flowering time. These differences appear robust to environmental variations as they are mostly preserved from one year to the other. They could be explained by random mutations appearing through the selection process and contributing or not to the response to selection.

### 4.3 Phyllochron temporal trends and climate

Phyllochron appears as significantly non-constant on almost all years and genotypes. Importantly, the model does not favor any temporal trend, thus the observed dynamics of phyllochron only originate from the data. Notably, phyllochron displays strong seasonal variations in 2015, a year with a peculiar climatic signature, and the dynamic trend is associated with the ancestral line, with an increasing and a decreasing phase for MBS genotypes and two increasing phases for F252. This suggests interactions between environmental and ancestral line as previously reported in phyllochron control (Wang et al., 2019). However, since different genotypes do not experience the same environment at the same developmental stage, these differences may originate either from selection or from differences in environment (Yu and Goh, 2019). Therefore, we proposed a model that includes longitudinal climatic variables and thus enables a calendar time analysis, assuming ancestral line-climate interactions.

Our results indicate that the climatic variables altogether enable to partly recover the strong phyllochron seasonal variations observed in 2015, but the model does not explain the more moderate variations. The strong correlations between climate variables, mostly preserved from one year to the other, would make the variable selection non-robust and the interpretation potentially misleading. Qualitatively, year 2015 was particularly dry and sunny from beginning of May to mid-July, which may explain the general increase of leaf appearance time over this period. The two-phase phyllochron observed in 2015 questions its commonly accepted linearity. This pattern may arise from a two-phase response to drought: a slowing leaf appearance allowing the plant to tolerate short term drought, followed by a recovering of the initial rate allowing the plant to complete its life cycle in case of long lasting drought conditions. Indeed, NeSmith et al. (NeSmith and Ritchie, 1992) induced such a response in corn using shelters to prevent rainfall during three weeks in 10-leaves plants. As mentioned above, the classic assumption of a constant phyllochron may prevent the ability to detect precisely these kinds of patterns with a slow-down followed by an acceleration. Furthermore, other physiological observations in corn during drought stresses suggested changes in leaf elongation rates, photosynthesis and transpiracy rates as well as leaf rolling that could results in changes of leaf appearance rate Markelz et al. (2011); Roth et al. (2013). The effect of irradiance on phyllochron has previously been investigated leading to different conclusions depending on the genotypes. While (Warrington and Kanemasu, 1983) did not detect any effect of irradiance on leaf emergence rate, (Birch et al., 1998) showed that decreased irradiance decelerates leaf emergence.

Out of the limitations of the two-step approach discussed above, additive modelling of covariate effects may not be enough to recover complex interplays between climate variables. Notably, the model does not account for potential non-monotonous effect of variables (medium optimum value). Nevertheless, a richer model would be at the price of a large number of parameters with respect to the number of phyllochron estimates used for model fitting, and could not be properly inferred with our limited number of genotype-year combinations. For instance, a study aiming at deciphering the non-additive effects of drought and temperature on crop yield required more than 50 years of climate variables (Matiu et al., 2017)

### 4.4 Conclusion

We developed a hypothesis-testing model for phyllochron based on a stochastic process, which combines a flexible and accurate modelling with an unbiased statistical testing procedure. The model enables to detect fine differences between related genotypes up to a moderate experimental effort (10 to 20 measurements throughout the season on 30-50 plants by genotypes). On the DSE dataset, we showed that the major sources of differences for the phyllochron were not the selection population (Early or Late), but rather the ancestral line (F252 or MBS), the year of experimentation, and the leaf rank. Moreover, our results clearly indicate that phyllochron is not always constant throughout the season, and these temporal trends could be associated with climate. These findings could be validated by implementing the phyllochron model with broader data sets (Millet et al., 2019).

## Author contributions

Methodology, software and visualization: S. Plancade and S. Huet contributed to the definition and estimation of the phyllochron model and the statistical framework and its numerical implementation; E. Marchadier, S. Plancade and C. Dillmann contributed to the definition and implementation of the climate model. Investigation: E. Marchadier, A. Ressayre and C. Dillmann participated to the definition of experimental protocol and to data collection and preparation. Writing: all authors discussed the results and contributed to the final manuscript. C. Dillmann supervised the ITEMAIZE project. Camille Noûs contributed to the collegial construction of the standards of science, by developing the methodological framework, the state-of-the-art, and by ensuring postpublication follow-up.

## Acknowledgements

This work was supported by a grant from the BASC labex (ITEMAIZE project) supported by IDEX Paris-Saclay. The authors declare no conflict of interest.

## Supplementary data

The file Supplementary-Materials.pdf provides technical details regarding mathematical methods and statistical analyses, together with secondary results and figures.

## Data availability statement

All scripts and data can be found at https://doi.org/10.15454/CUEHO6

## Supplementary material

### A Monte Carlo Expectation Maximisation Algorithm

Consider a given genotype *lsg* (or higher level genotypic group) and year *y*, the indexes *lsg* and *y* are omitted in this section. For a plant *p*, let (*t*_*p*,1_,…, *t_p,N_p__*) be the monitoring times (with time origin at sowing) and *X_p,j_* the number of leaves of plant *p* at time *t_p,j_*. For each plant and leaf rank, the observations indicate that leaf *f* appeared at some time between the last observation date when the plant had at most (*f* – 1) leaves, and the first observation date when the plants had at least *f* leaves. We denote by *ν_p,f_* and *τ_p,f_* respectively these two dates:

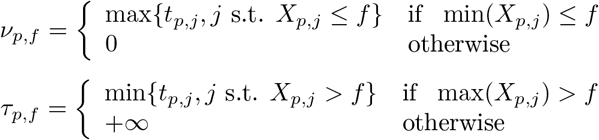

The likelihood of the observations involves an integral of dimension *f*_max_ – *f*_min_ + 1, thus a direct maximisation would be intractable. Thus, we considered a Monte-Carlo Expectation Maximisation algorithm, wihere the *complete data* (unobserved) are **H**_p_ = (*H_p,f_*)_*p*=1,…, *n,f*=*f*_min_,…, *f*_max__

- <u>Initialisation</u>. Proxies of the {*H_p,f_*}*_p,f_* are computed as:

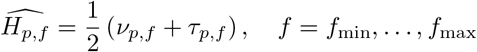

for all (*p, f*) such that *ν_p,f_* > 0 and *τ_p,f_* < ∞, and the initial values of *μ_f_* and *σ_f_* are inferred as robust estimators of mean and variance based on the proxies.
- <u>Monte-Carlo E-step</u>. Let 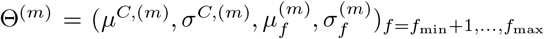 be the current value of the parameters. For each *p* a sample 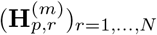 is generated from the conditional distribution:

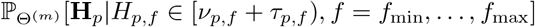

namely a truncated multivariate Gaussian distribution. Computation time of classic rejection methods dramatically increase with the dimension, nevertheless specific are available for the truncated multivariate distribution. We used the R package `TruncatedNormal` (Botev and Belzile, 2020) well adapted when the truncated region is in the tail of the distribution.
- <u>M-step</u>. For *f* = *f*_min_ + 1,…, *f*_max_, let 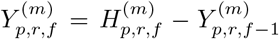. The new value of the parameters Θ^(*m*+1)^ maximises

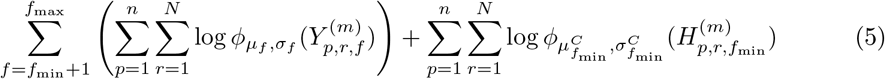

with *φ_μ,σ_* the density of the normal distribution with mean *μ* and standard deviation *σ*. This problem is equivalent to a simple maximum likelihood estimation of a multivariate Gaussian distribution with diagonal covariance matrix, thus:

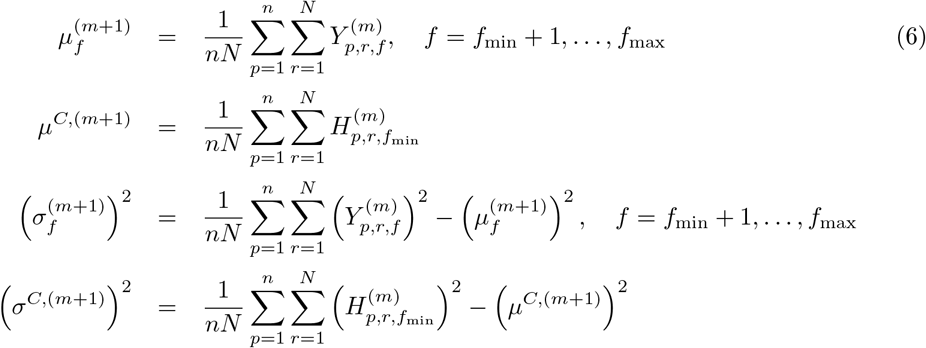 The algorithm is iterated until stabilisation of the parameters.

### B Parametric sub-models

For each genotype lsg and year y, we consider the following parametric models for the average interval times between leaves 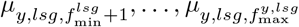 as a function of leaf rank:

- Constant: *μ_y,lsg,f_* = *a_y,lsg_*, 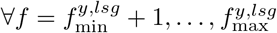 (mod-ct)
- Linear: *μ_y,lsg,f_* = *a_y,lsg_* × *f* + *b_y,lsg_*, 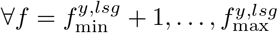 (mod-lin)
- Piecewise constant: 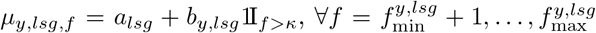 for *κ* = *f*_min_ + 2,…, *f*_max_ – 2 (mod-pct(*κ*))
- Piecewise linear: 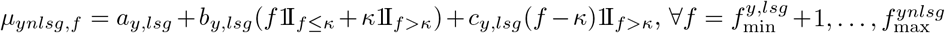 for *κ* = *f*_min_ + 3,…, *f*_max_ – 3 (mod-plin(*κ*))

In order to test the classic hypothesis of constant leaf appearance rate which corresponds to (mod-ct), for each genotype we compared models (mod-lin), (mod-pct(*κ*))), (mod-plin(*κ*)) with (mod-ct). First, (mod-ct) was compared to all parametric models and to the complete model using the *χ*^2^-likelihood ratio test; if all *p* > 0.01, (mod-ct) is selected; otherwise, the parametric model with the largest AIC is selected.

#### Monte Carlo EM algorithm for the parametric sub-models

We omit the indexes *y* and *lsg*. The parametric models we considered are equivalent to imposing a linear constraint:

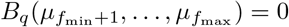

with *B_q_* a *q* × *s*-matrix, *s* = *f*_max_ – *f*_min_ and *q* < s. The parameter *μ^C^* is unconstrained.

- Constant model (mod-ct):

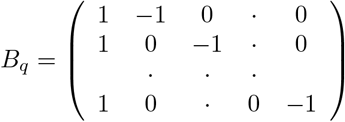
- Linear model: *B_q_* is the (*s* – 2) × *s* matrix such that for every *j* > 2

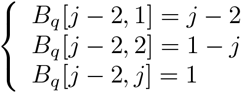

and the other coefficients are zero.
- Piecewise constant (mod-pct(*κ*)): let *κ′* = *κ* – *f*_min_, then *B_q_* is the (*s* – 2) × s matrix with,

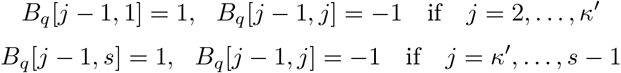

and the other coefficients are zero.
- Piecewise linear: let *κ′* = *κ* – *f*_min_ *B_q_* is the (*s* – 3) × *s* matrix such that for every 2 < *j* ≤ *κ′*

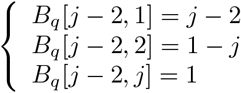

and for every *κ′* + 1 < *j* ≤ *s*

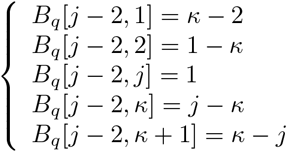

and the other coefficients are zero.

**Table S1:**
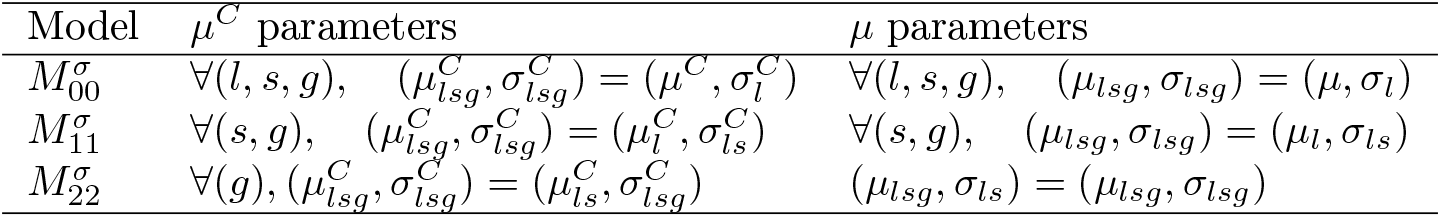
Models in which the mean parameters depend on grouping level *j* – 1 and the variance parameters on grouping level *j*

The Monte-Carlo E-step is identical to the case of unconstrained *μ*. The M-step corresponds to the maximum likelihood estimation problem from a multivariate Gaussian distributed sample under linear constraints on the mean, which admits an explicit expression as follows (Zoppé et al., 2001). Let *B_s-q_* be any matrix such that

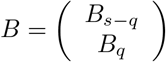

is invertible. Then, let

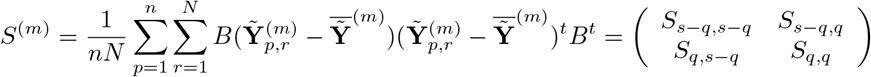

where 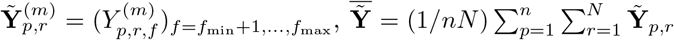, and *S_a,b_* are submatrices of *S* with dimension *a* × *b*. Let 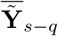 (resp. 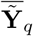) be the subvector of 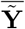 with the *s – q* first (resp. the *q* last) coordinates of 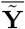, then

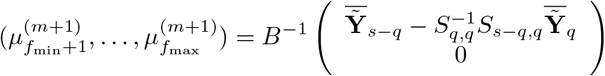

### C Are genotypic group effect due to differences in the average phyllochron?

The differences of phyllochron between genotypic groups (genotypes, selection populations and inbred lines) could originate from differences in the average phyllochron (*μ^C^, μ*) and/or on the standard deviations (*σ^C^,σ*). In this subsection, we perform additional comparisons to ensure that the genotypic effect indeed impacts the average phyllochron. In addition to models *M*_*j*–1,*j*–1_ and *M_j,j_* in Table 3, we considered models 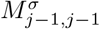 in which the mean parameters depend on grouping level *j* – 1 and the variance parameters on grouping level *j* (Table S1)

The comparison of models *M_j,j_* and 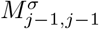 enables to test the effect of genotypic grouping on the mean parameters (*μ_y,lsg,f_*) only, regardless of its effect on variance parameters (*σ_y,lsg,f_*). Results in Table S2 indicate that these comparisons lead to similar p-values than the comparison *M_j,j_/M*_*j*–1,*j*–1_. Moreover, the estimates of (*μ_y,lsg,f_*) computed under models *M*_*j*–1,*j*–1_ and 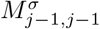 are very similar (Figure S2). Therefore, the observed genotypic groups effects are due, in a significant part, to differences on the average phyllochron 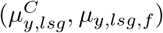 rather than on the nuisance parameters 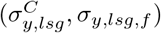.

### D Criteria for model comparison - Evaluation through permutation test

Intially, we considered three criteria for model comparisons.

- the AIC criterion –2*ℓ* + 2*p* with *ℓ* the log-likelihood of the observations and *p* the number of parameters in the model. The smaller the better.
- the BIC criterion –2*ℓ* + *p* × log *n* with *n* the sample size (number of plants). The smaller the better. As soon as the sample size is larger than eight, BIC is more conservative than AIC.
- The p-value of the *χ*^2^-likelihood ratio test.

These three criteria rely on asymptotic heuristics, and there is no obvious choice a priori. On the contrary, permutation tests are unbiased but too time consuming to be used for all comparisons. Thus, in order to evaluate the relevance of the three criteria for finite sample size, we used a permutation test with N = 200 permutations for the comparison *M*_22_/*M*_33_ for the two genotypes of the group F252-Early. On the non-permuted data, AIC selected *H*_1_ (1904 *versus* 1915) indicating a genotypic effect, while BIC (which is more conservative by definition) selected *H*_0_ (1954 *vs* 1982), indicating no genotypic effect. The *χ*^2^-likelihood ratio test had a significant p-value (5.10^-5^), leading to the same conclusion that the AIC criterion. Therefore, we wondered if BIC criterion is too conservative, or if AIC and *χ*^2^-test over-estimated the genotypic effect.

On the one hand, the unbiased permutation test had a p-value of 0.02, coherent with the *χ*^2^-log likelihood test, even less significant. Besides, the quantile 0.05 of the empirical distribution of the *χ^2^* p-value on the resampled data, equal to 0.013, could be considered as a conservative heuristic threshold, since tests involving more plants (e.g. selection or line effect) are expected to be less biased. Nevertheless, given the very weak p-values obtained on the data, this distinction would not dramatically change the conclusions. On the other hand, the AIC criterion proved to be reliable since it mostly selects *M*_22_ on the resampled data (94.5% of the permutations), even slightly too selective; similarly to *χ*^2^-likelihood test. Finally, the BIC criterion appeared too conservative, as it did not allowed to detect a significant genotypic effect with the original genotypes.

**Figure S1:**
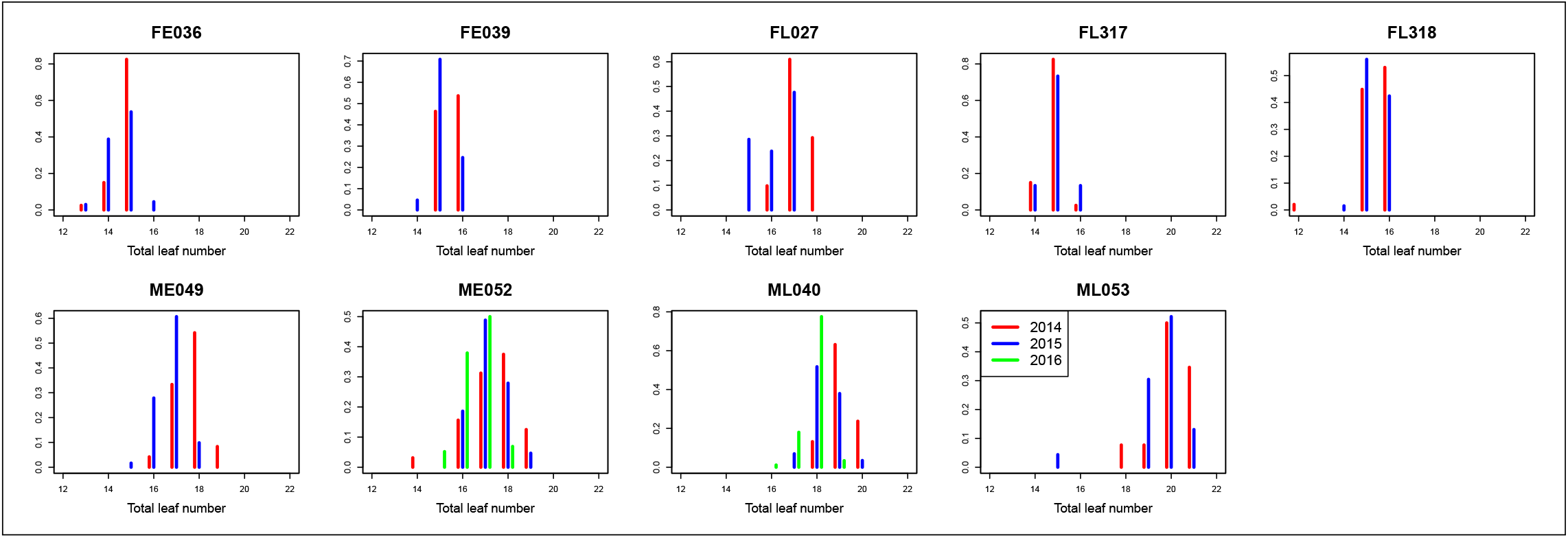
Distribution of the total leaf number for all plants by genotype and by year.

**Figure S2:**
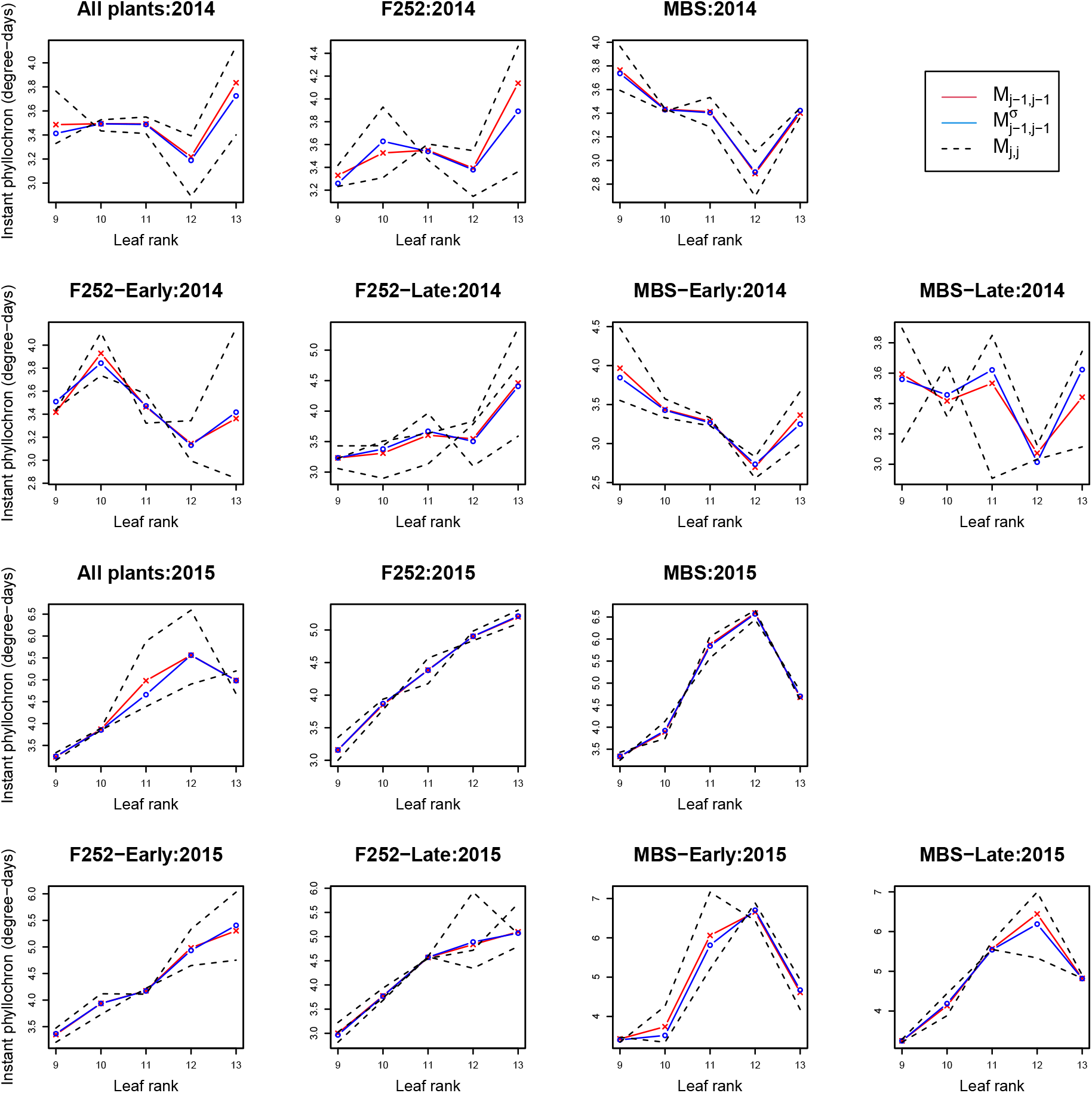
Estimates of instant phyllochron for models *M*_*j*–1,*j*–1_ ((*μ^C^, μ, σ^C^, σ*) depend on the grouping level *j* – 1, red), *M_j,j_* ((*μ^C^, μ, σ^C^, σ*) depend on the grouping level *j*, dashed lines) and 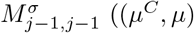 ((*μ^C^, μ*) depend on the grouping level *j* – 1 and (*σ^C^, σ*) depend on the grouping level *j*, blue). Models are described in Tables 3 and S1. Rows 1 and 2 correspond to year 2014, and rows 3 and 4 to 2015. For each year, plots entitled All plants correspond to *j* = 1, plots F252 and MBS to *j* = 2, and plots F252-Early, F252-Late, MBS-Early and MBS-Late to *j* = 3.

**Table S2:** *χ*^2^ likelihood ratio p-value for genotypic group effect on the mean and variance of the phyllochron (*M_j,j_*/*M*_*j*–1,*j*–1_) and on the average phyllochron μ only 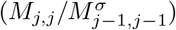.

| 2014 |  |  | 2015 |  |  |
| --- | --- | --- | --- | --- | --- |
| | $M_{j,j}/M_{j-1,j-1}$ | $M_{j,j}/M_{j-1,j-1}^\sigma$ | | $M_{j,j}/M_{j-1,j-1}$ | $M_{j,j}/M_{j-1,j-1}^\sigma$ |
| all | <e-16 | 7.3e-15 | all | <e-16 | <e-16 |
| F | 2.00e-12 | 1.8e-09 | F | 3.0e-03 | 1.7e-02 |
| M | 4.20e-02 | 7.2e-02 | M | 1.3e-05 | 2.0e-04 |
| Fearly | 6.60e-05 | 3.6e-05 | Fearly | 4.9e-04 | 6.8e-05 |
| Flate | 5.70e-12 | 1.1e-9 | Flate | 3.7e-09 | 9.4e-06 |
| Mearly | 6.00e-02 | 7.3e-02 | Mearly | <e-16 | <e-16 |
| Mlate | 3.80e-05 | 6.3e-06 | Mlate | 1.5e-04 | 2.0e-05 |

**Figure S3:**
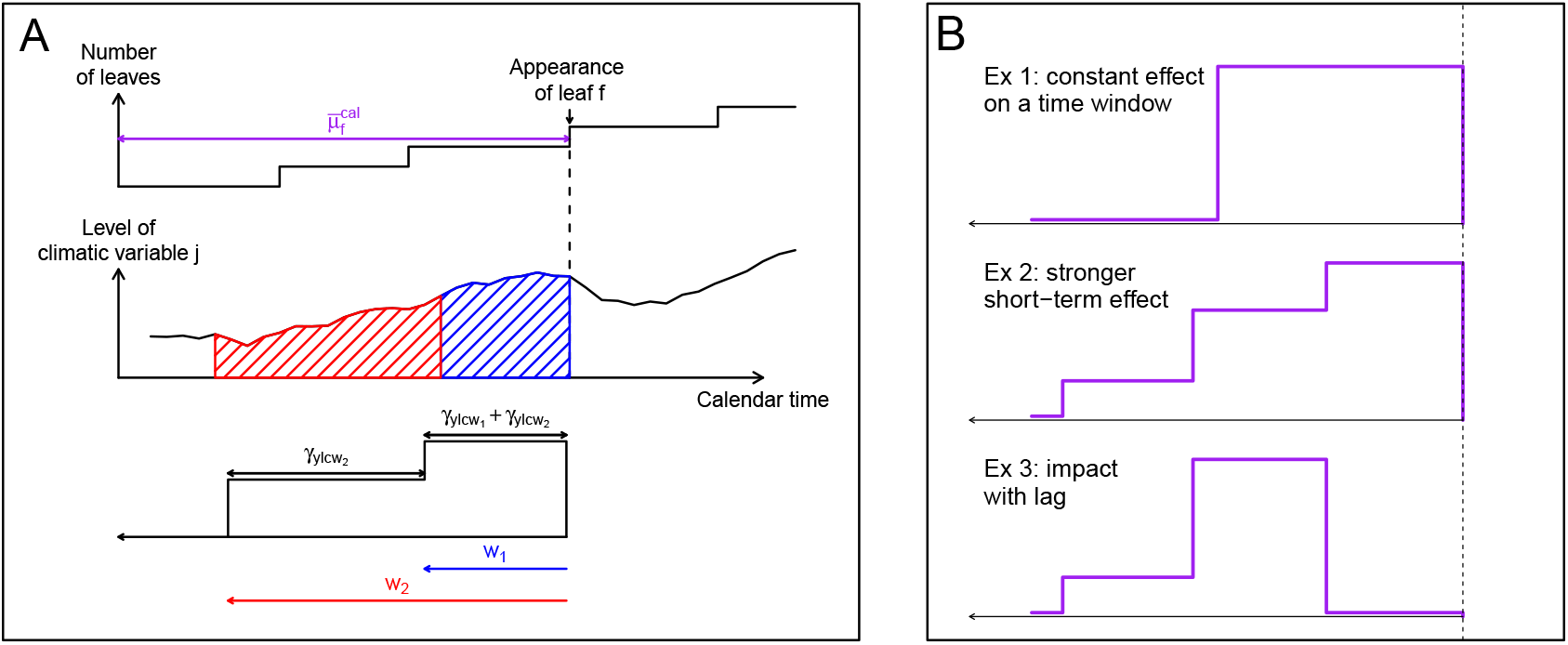
Climatic model. (A) The graph at the top displays the average phyllochron process for a given genotype and year in calendar time; The graph below gives the daily value of a climatic variable; the piecewise constant function at the bottom corresponds to the weights applied to the climatic variable values (cf model (4)). On this example the vector (*γ_y,l,c,w_*)*_w_* has two non-zero coefficients for *w* = *w*_1_ and *w* = *w*_2_, therefore the impact of the climatic variable on *μ_y,lsg,f_* is equal to the sum of the daily values of the climatic variables on the time interval 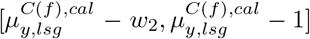 weighted by *γ*_*y,l,c,w*_1__) + *γ*_*y,l,c,w*_2__) on 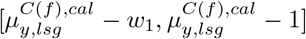 and by *γ*_*y,l,c,w*_2__) on 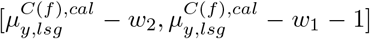. (B) Examples of behaviours that can be modelled by (4) with three non-zero coefficients in (*γ_y,l,c,w_*)*_w_*. In the first example, the time between appearance of successive leaves is assumed to result from the value of the climatic variable on a certain time window before appearance of all leaves, with the same impact on the whole window. On the second example, the impact of the covariate is assumed to be stronger for the value close to the leaf appearance. In the last example, the impact of the climatic variable is assumed to be subjected to a lag.

**Figure S4:**
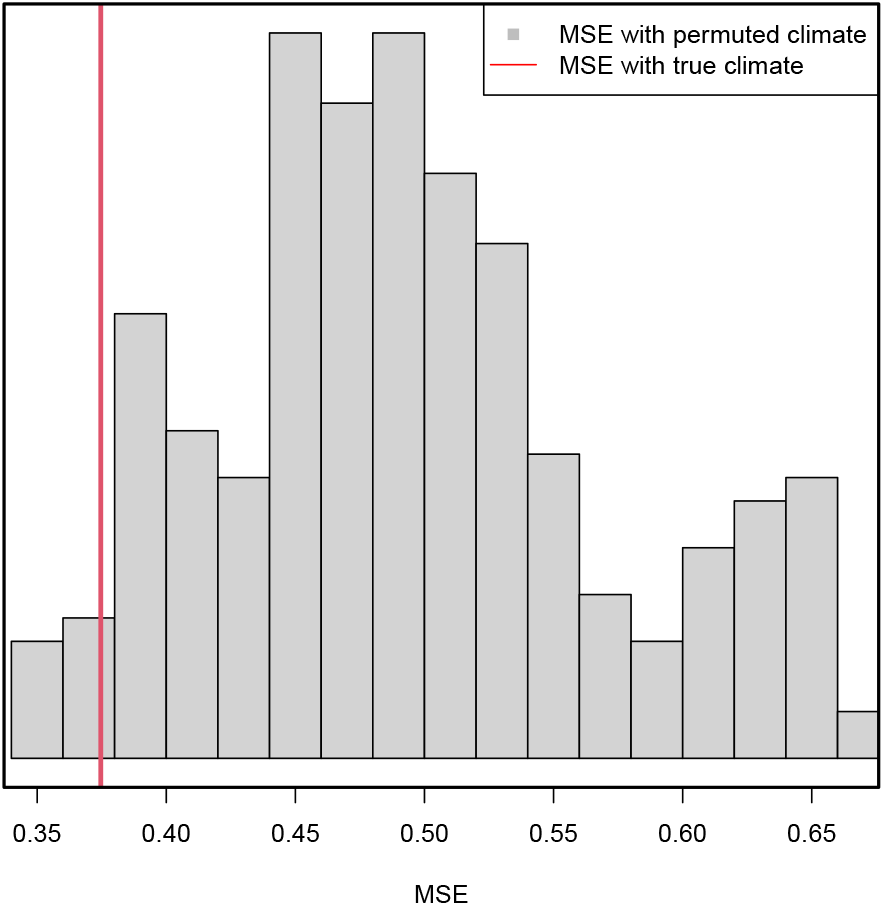
Prediction mean square error for the climatic model with false climates. Histogram of the prediction MSE for the climatic model with 252 false climate and prediction MSE with the true climates (red line).

**Figure S5:**
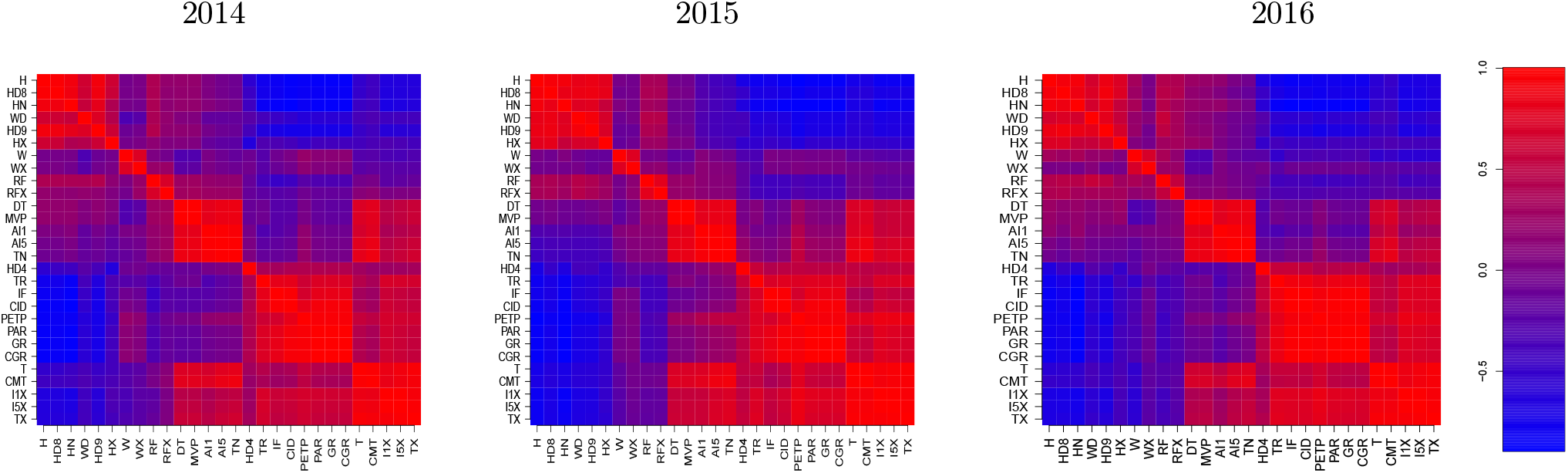
Pearson correlation between climatic variables on year 2014,2015 and 2016. Orders of variables in rows and columns is identical for the 3 years. Climatic variables are described in Table 1.

**Figure S6:**
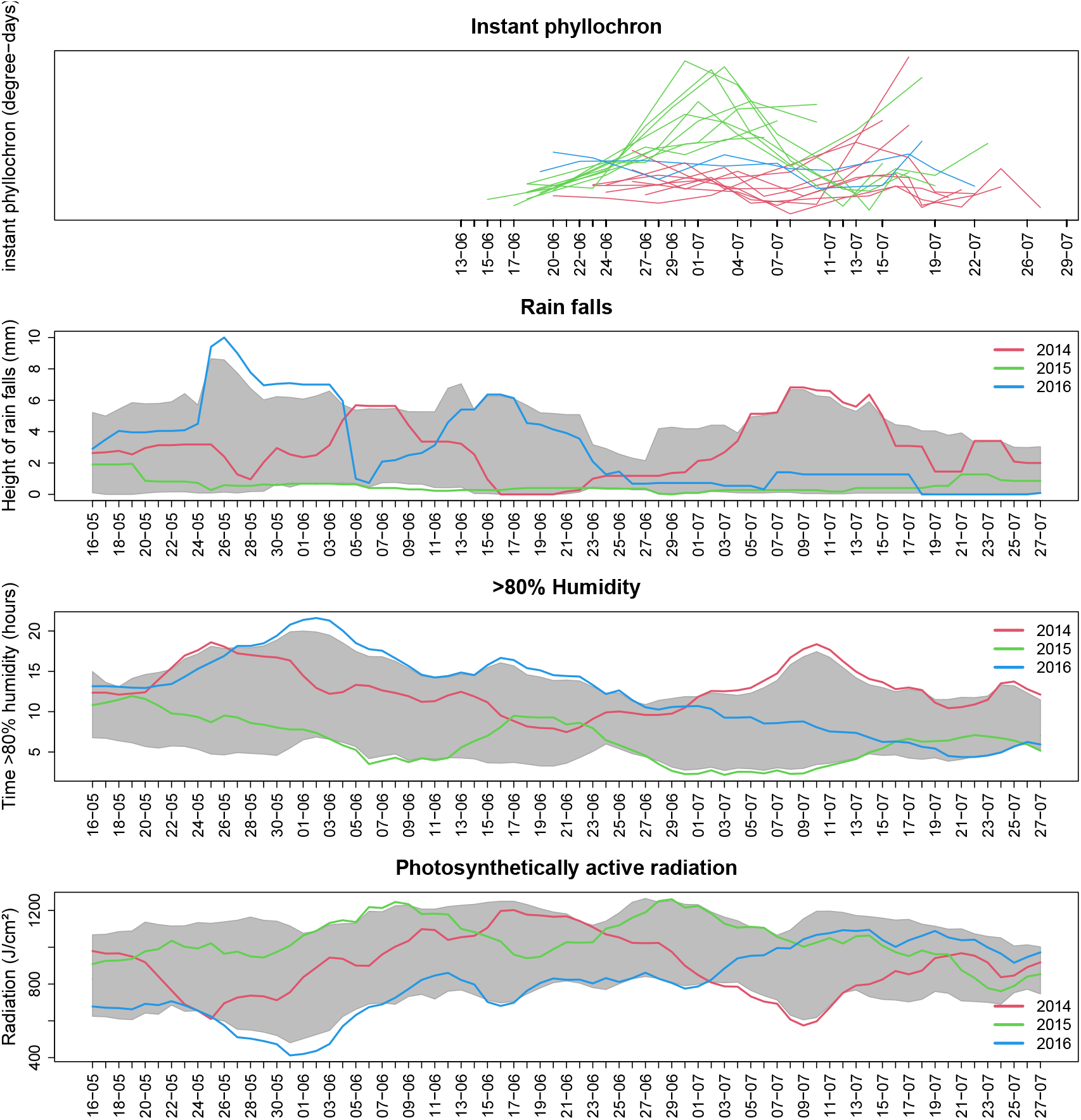
Instant phyllochron and climate variables recorded in Saclay for May-July period from 2009 to 2017. The higher panel show the instant phyllochron of each of the genotypes during the 3 years of experimentation in function of calendar time. The following panels show the climate trends for the 2009-2017 period, as well as the specific curves for the years 2014, 2015 and 2016 for three variables related to rainfalls, humidity and photosynthetic radiations. Rain falls, time with >80% humidity and photosynthetically active radiation are daily recorded and smoothed on a −5/+5days window. Grey area represents the intervals encompassing the daily climate variables of 7 out of the 9 years (2009 to 2017) Patterns of 2014, 2015 and 2016 are indicated in colors (respectively red, green and blue)

